# Uncertainty-aware genomic deep learning with knowledge distillation

**DOI:** 10.1101/2024.11.13.623485

**Authors:** Jessica Zhou, Kaeli Rizzo, Trevor Christensen, Ziqi Tang, Peter K Koo

**Affiliations:** Simons Center for Quantitative Biology, Cold Spring Harbor Laboratory, NY, USA; Currently at InstaDeep, New York, NY, USA

## Abstract

Deep neural networks (DNNs) have advanced predictive modeling for regulatory genomics, but challenges remain in ensuring the reliability of their predictions and understanding the key factors behind their decision making. Here we introduce DEGU (Distilling Ensembles for Genomic Uncertainty-aware models), a method that integrates ensemble learning and knowledge distillation to improve the robustness and explainability of DNN predictions. DEGU distills the predictions of an ensemble of DNNs into a single model, capturing both the average of the ensemble’s predictions and the variability across them, with the latter representing epistemic (or model-based) uncertainty. DEGU also includes an optional auxiliary task to estimate aleatoric, or data-based, uncertainty by modeling variability across experimental replicates. By applying DEGU across various functional genomic prediction tasks, we demonstrate that DEGU-trained models inherit the performance benefits of ensembles in a single model, with improved generalization to out-of-distribution sequences and more consistent explanations of cis-regulatory mechanisms through attribution analysis. Moreover, DEGU-trained models provide calibrated uncertainty estimates, with conformal prediction offering coverage guarantees under minimal assumptions. Overall, DEGU paves the way for robust and trustworthy applications of deep learning in genomics research.

---

Deep neural networks (DNNs) have demonstrated strong performance in predicting the outputs of functional genomics experiments directly from DNA sequences [1–3]. By approximating experimental assays, these DNNs enable virtual experiments that explore the functional effects of genomic sequence perturbations. Applications include predicting the impact of genetic variants [1, 4, 5], simulating CRISPR-like perturbation experiments that uncover regulatory rules of sequence motif syntax [6–8], and designing novel DNA sequences [9–14]. In these applications, high-performing DNNs serve as black-box *in silico* oracles or scoring functions, mapping DNA sequence inputs to a target molecular phenotype, such as gene expression or chromatin accessibility. These models have the potential to improve hypothesis generation and guide more optimal experimental design, setting the stage for efficient AI-guided biological discovery.

However, these downstream applications assume that DNNs maintain their predictive performance even when the statistical properties of the input data differ from those seen during training, a phenomenon known as a covariate shift [15]. Model generalization is often assessed using in-distribution held-out sequences from the same experiment that generated the training data. While these held-out sequences come from different genomic regions, they are typically similar in genomic composition to the training data due to evolutionary constraints. Consequently, this assessment may not accurately reflect the model’s ability to predict sequences with greater genetic variation, beyond the natural genome.

Recent work has shown that state-of-the-art genomic DNNs, such as Enformer [1], generalize well to predicting single-nucleotide variant effects within cis-regulatory elements [16, 17]. However, these predictions are typically provided without calibrated measures of uncertainty, making it difficult to determine when a prediction can be trusted. The lack of reliable uncertainty estimates limits the utility of genomic DNNs in downstream applications such as disease variant interpretation. Thus, developing methods to quantify and improve uncertainty calibration represents a critical challenge for advancing variant effect prediction.[18, 19].

One approach to quantifying uncertainty involves training an ensemble of DNNs, where each model typically shares the same architecture but differs in its randomly initialized parameters, allowing the variation across their predictions to serve as an empirical measure of confidence [20]. These so-called *deep ensembles* [21] improve performance by averaging predictions across diverse models, where each model captures a different aspect of the data. Averaging reduces individual model errors and balances biases, leading to more accurate and robust predictions than any single model alone. Ensembling has indeed proven to be an effective strategy to improve predictive performance for genomic DNNs [22–25].

In addition to improving prediction accuracy, deep ensembles are valuable for quantifying epistemic uncertainty, or model uncertainty due to limited data, through the variability across the ensemble predictions [21]. For example, a recent study leveraged deep ensembles of genomic DNNs to investigate relationships between small sequence perturbations (e.g., expression quantitative-trait loci) and uncertainty in predictions [26]. Deep ensembles also provide more reliable post hoc explanations when averaging the attribution maps from each model in the ensemble. These attribution maps assign importance scores to nucleotides in a given sequence, revealing sequence motifs that are functionally relevant for the model’s predictions [17, 27–29]. Despite these advantages, deep ensembles face several challenges that limit their practicality in genomics applications. One major issue is the increased computational overhead required to train and deploy multiple models, making large-scale inference tasks such as genome-wide variant effect predictions or extensive *in silico* experiments computationally expensive. Additionally, while deep ensembles capture epistemic uncertainty, they fail to account for aleatoric uncertainty [30, 31], the irreducible noise stemming from technical and biological variability inherent in biological data. Consequently, deep ensembles may provide an incomplete measure of total predictive uncertainty. Furthermore, the need to manage and maintain multiple models adds substantial complexity to implementation, creating scalability challenges, especially as model architectures continue to trend towards greater size and complexity [1, 24, 32–36].

To address these limitations, we introduce DEGU (Distilling Ensembles for Genomic Uncertainty-aware models), a method that combines ensemble learning and knowledge distillation [37] to improve the robustness and explainability of DNN predictions. DEGU leverages ensemble distribution distillation [38], a variant of knowledge distillation that focuses on learning the distribution of predictions from the ensemble rather than individual point estimates. This is accomplished by training a single student model in a multitask fashion to perform two primary tasks: 1) predict the mean of the ensemble’s predictions, and estimate the corresponding epistemic uncertainty based on the variability across the ensemble’s predictions. Additionally, DEGU can incorporate an optional auxiliary task for the student model to predict the aleatoric uncertainty, estimated from the variability observed across experimental replicates.

By applying DEGU to different DNNs across various functional genomics prediction tasks, we found that distilled models exhibit improved generalization and enhanced robustness in their attribution maps compared to standard training methods. Furthermore, DEGU-distilled models accurately predict epistemic uncertainty. Together, DEGU provides the efficiency of a single model during inference while preserving the performance and robustness of deep ensembles, with the added benefit of generating calibrated uncertainty estimates.

## Results

### DEGU: distilling the knowledge of ensembles into a single model

DEGU employs ensemble distribution distillation [38] to transfer the collective knowledge from an ensemble of models, which we refer to as teacher models, to a single, student model (Fig. 1). The process begins with the creation of a teacher ensemble, where multiple DNNs are trained independently with different random initializations. Through knowledge distillation, the student model learns the distribution of predictions from the teacher ensemble by performing multiple tasks concurrently: 1) predicting the mean of the ensemble’s predictions and 2) predicting the variability across the ensemble’s predictions. This assumes the predictions across the teacher ensemble follow a normal distribution, characterized by a mean, which is the standard quantity used to distill an ensemble’s predictions, and standard deviation, which reflects epistemic uncertainty. Additionally, when at least three experimental replicates are available, the student model can optionally be trained to predict the aleatoric uncertainty [39], which is approximated by the variability observed across replicates in the training data. Our multitask learning approach ensures that DEGU captures the distribution of predictions from the ensemble along with the variability inherent in the data. This enables DEGU to retain the performance and robustness advantages of deep ensembles while reducing computational overhead that scales proportionally to the ensemble size (by 90% in our case) during downstream inference tasks (Supplementary Table 1).

**Figure 1.**
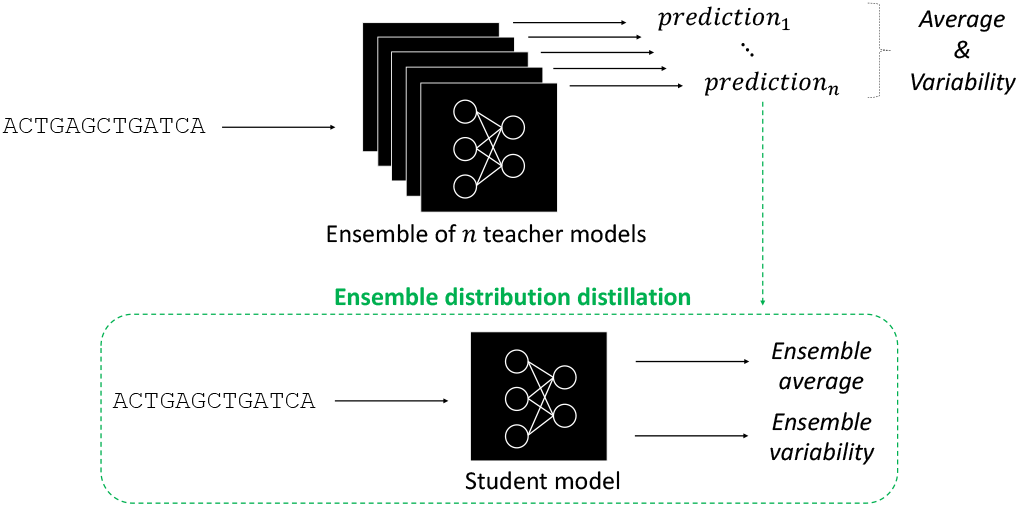
Schematic of ensemble distribution distillation with DEGU. For a given input, the predictions from an ensemble of teacher models are used to generate target labels for training a distilled student model. The student model is trained to predict both the average and variability of the predictions from the teacher ensemble.

### DEGU approximates the predictive performance of deep ensembles

To demonstrate DEGU’s utility, we applied it to various DNNs trained on diverse datasets: fly enhancer activity measured via STARR-seq for developmental (Dev) and housekeeping (Hk) promoters [40], human cis-regulatory sequence activity quantified through lentiMPRA for K562 and HepG2 cells [23], and base-resolution ATAC-seq profiles from a human cell line [41]. For each application, we constructed a teacher ensemble consisting of 10 models using established architectures suited to each task: multi-task DeepSTARR [40] for fly enhancers, single-task ResidualBind [6, 42] for lentiMPRA [23], and a standard convolutional neural network (CNN) for base-resolution ATAC-seq profiles [8]. Student DNNs were then trained with DEGU’s ensemble distribution distillation procedure, using the same architecture as their respective teacher models (see Methods).

To simulate scenarios with limited data, we compared different data size regimes by randomly downsampling the original training data while retaining the full validation and test sets for model evaluation. Strikingly, the distilled DNNs (i.e., the student models) outperformed DNNs with standard training (Fig. 2), approaching the performance of the teacher ensemble. Although the student models shared the same architecture as the teacher, training with ensemble-generated target labels consistently improved predictive performance and often matched the performance of ensembles consisting of 10 models, particularly in low-data regimes. For example, distilling DeepSTARR with only 25% of the STARR-seq training data yielded performance comparable to standard training on the full dataset for both developmental and housekeeping promoters (Fig. 2a-b). Similar gains were observed on lentiMPRA data, with stronger improvements in K562 and more modest but noticeable effects in HepG2. For base-resolution ATAC-seq profiles, distilled models achieved performance comparable to standard training, with only a slight reduction relative to teachers (Supplementary Fig. 1a). Next, we extended DEGU to DREAM-RNN models [43], which have previously demonstrated state-of-the-art accuracy on the DeepSTARR and lentiMPRA datasets (Supplementary Fig. 2). Distilled DREAM-RNN models trained on lentiMPRA data closely tracked the ensemble, while improvements on DeepSTARR were more modest and most evident as training data increased.

**Figure 2.**
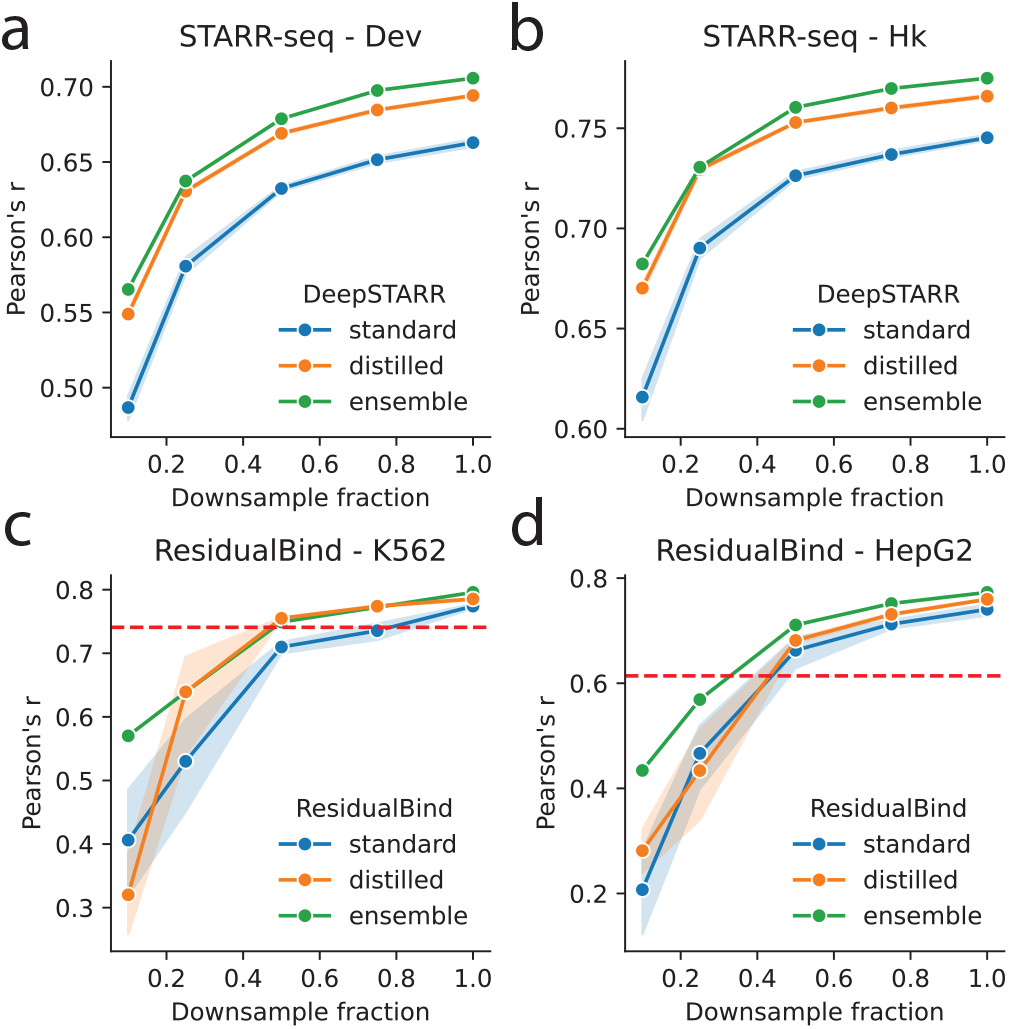
Comparison of model performance with downsampled training data. (**a**,**b**) Predictive performance of DeepSTARR models trained on different subsets of randomly downsampled STARR-seq data for (**a**) developmental (Dev) and (**b**) housekeeping (Hk) promoters. (**c**,**d**) Predictive performance of ResidualBind models trained on subsets of randomly downsampled lentiMPRA data for(**c**) K562 and (**d**) HepG2 cell lines. (**a**-**d**) Plots include models with standard training (blue; *n* = 10), DEGU-distillation (orange; *n* = 10); and the ensemble average of the teacher models with standard training (green). Shaded regions indicate 95% confidence intervals. Red dashed lines in (**c**,**d**) correspond to MPRAnn performance trained on the full lentiMPRA dataset for corresponding cell type.

We also observed additional performance gains when the performance of individual models within the teacher ensemble were improved (Supplementary Fig. 3). In this scenario, every model (including the teacher and student) was trained with evolution-inspired data augmentations provided by EvoAug [44, 45]. This resulted in higher performance gains for the individual models, the ensemble, as well as the distilled models (Supplementary Fig. 3). Moreover, as the number of models in the teacher ensemble increased, the ensemble’s predictive performance improved and plateaued around *n* = 10 models. The performance of the distilled models also plateaued but with smaller ensembles of around *n* = 5 models (Supplementary Fig. 4). Notably, the performance gap between the ensemble and the distilled models widened as the ensemble size increased beyond this range.

### DEGU improves generalization under covariate shifts

Most downstream applications require genomic DNNs to generalize well under covariate shifts, especially when making predictions for sequence perturbations that were not represented in the training data. Ensembles are typically expected to improve out-of-distribution (OOD) generalization [21, 46] because they aggregate variable predictions, thereby smoothing out arbitrary behavior in regions with limited or no data (Fig. 3a). Therefore, we hypothesized that DEGU-distilled models, which approximate the ensemble’s function, would also generalize better to OOD sequences. However, systematically assessing OOD generalization is challenging due to the limited availability of appropriate OOD data; that is, experimental measurements in the same biological system for sequences with matched levels of genetic variability as the downstream application task. To circumvent this issue, we instead used a proxy for OOD generalization by evaluating how closely the distilled models approximated the teacher ensemble’s behavior under varying levels of simulated covariate shift. Specifically, we created three new sets of test sequences based on the original STARR-seq test sequences, each simulating a different degree of distribution shift: (1) partial random mutagenesis at a rate of 0.05 at each position of the sequence to introduce a small shift, (2) evolution-inspired mutagenesis provided by EvoAug [44, 45] to create an intermediate shift, and (3) randomly shuffled sequences to represent a large shift (see Methods). The small shift introduced by partial random mutagenesis likely preserved most key motifs. In contrast, the intermediate shift, generated through evolution-inspired mutagenesis created more substantial compositional rearrangements in regulatory sequences. The large shift, created by randomly shuffling the sequences, likely disrupted and inactivated many functional regions, resulting in lower predicted regulatory activity overall (Supplementary Fig. 5).

**Figure 3.**
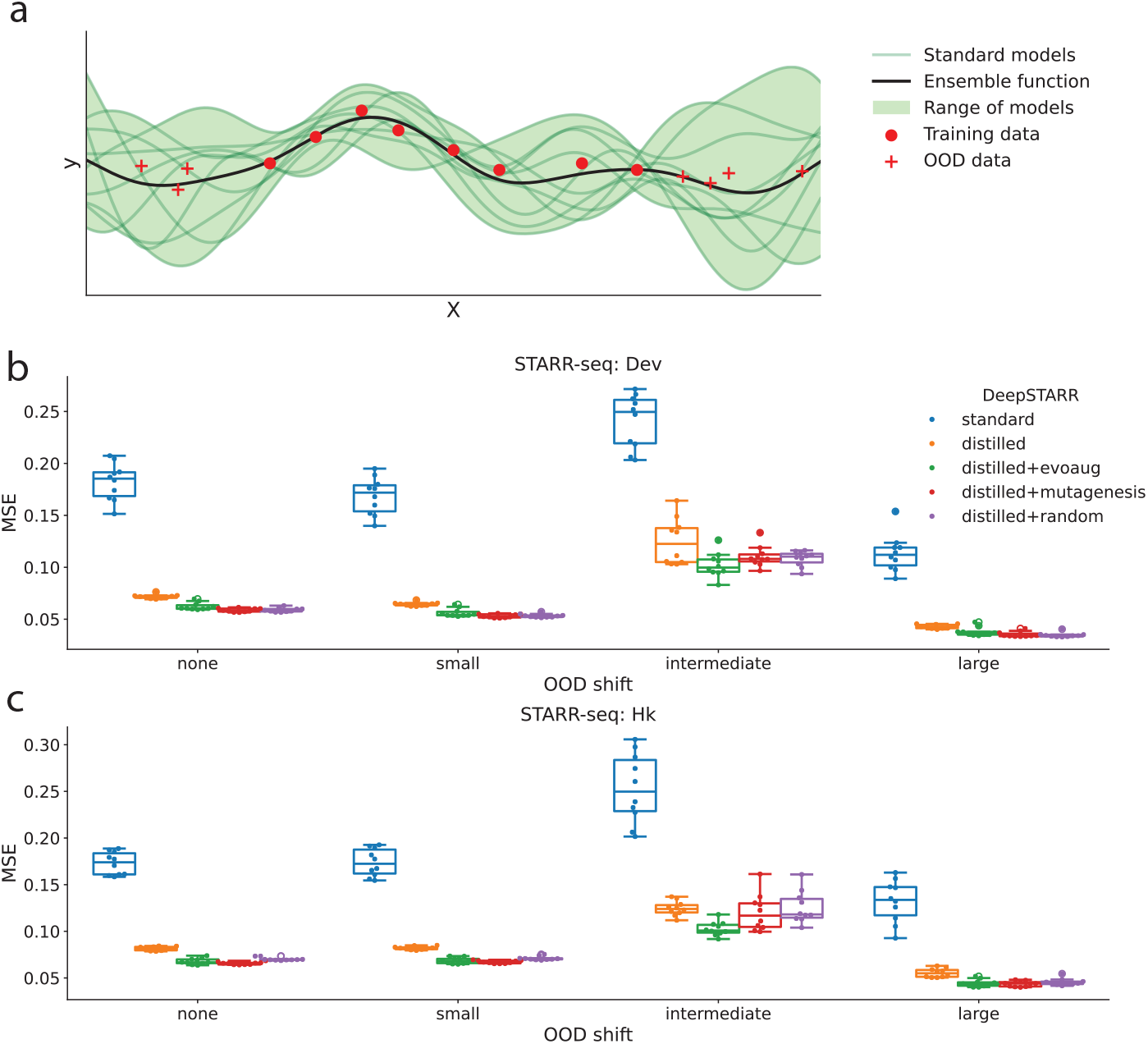
DeepSTARR OOD generalization performance. (**a**) Toy example illustrating 1-dimensional functions fitted by individual models trained with different random initializations (green lines) on training data (red points), along with the ensemble average function (black line). Sampling data in OOD regions, with labels provided by the teacher ensemble (red plus signs), stabilizes the distilled model’s function approximation in OOD regions. (**b**,**c**) Mean squared error (MSE) between the teacher ensemble (DeepSTARR trained with EvoAug) and individual DeepSTARR models for (**b**) developmental (Dev) and (**c**) housekeeping (HK) promoter activity across different model training procedures: standard training (blue), DEGU-distillation (orange), DEGU-distillation with dynamic EvoAug mutagenesis (green), DEGU-distillation with dynamic partial random mutagenesis (red), and DEGU-distillation with dynamic randomly shuffled sequences (purple). Results are shown for sequences with varying degrees of distribution shift: none (original test set), small (partial random mutagenesis), intermediate (EvoAug mutagenesis), and large (random shuffle). Boxplots represent *n* = 10 models trained with different random initializations, with the boxes indicating the first and third quartiles, the central line indicates the median, and whiskers denote the data range.

We then calculated the mean-squared error (MSE) between each model’s predictions and an ensemble average of *n* = 10 EvoAug-trained DeepSTARR models, which we treat as an in silico oracle. As expected, the distilled DeepSTARR models provided consistently better approximations of the teacher ensemble compared to standard-trained DeepSTARR models (Fig. 3b-c). Interestingly, applying a large covariate shift via randomly shuffling the sequences resulted in lower MSE values, possibly due to the lower overall activity levels in these sequences (Supplementary Fig. 5).

Next, we explored whether exposure to OOD data during training could improve model generalization [47–49]. Specifically, we hypothesized that introducing OOD sequences with ensemble-generated labels would help the model better approximate the ensemble function in regions where training data is sparse. Without these additional samples, the model’s behavior in such regions could be arbitrary due to lack of training data. To test this, we applied the same transformations used to simulate covariate shifts in the previous analysis to each minibatch of sequences during training and used the teacher ensemble to generate corresponding labels for these transformed sequences. The augmented sequences and their new target values replaced the original STARR-seq training data, allowing us to train distilled DeepSTARR models with a more diverse set of training data. Training distilled student models with dynamic data augmentations provided consistent, though modest, performance gains on the original test set (Supplementary Fig. 6). Furthermore, these augmentations improved the student models’ function approximation to the teacher ensemble under different levels of covariate shift, with slightly better performance when the augmentations closely matched the target covariate shift level (Fig. 3b-c). These findings highlight the importance of incorporating training data beyond the natural genome to improve model reliability in downstream applications that require robust generalization covariate shifts.

### DEGU improves attribution analysis

Attribution methods assign an importance score to each nucleotide in a sequence, indicating how much that nucleotide contributes to the model’s prediction (as with DeepSHAP [50] and DeepLIFT [51]) or how sensitive the model’s output is to changes at that nucleotide (as with gradient-based Saliency Maps [52]). Visualizing attribution scores as a sequence logo can reveal biologically meaningful patterns, such as transcription factor binding motifs [7, 40, 53]. However, attribution methods can be sensitive to local variations in the model’s learned function, which can arise when fitting to noise in the data. This variability may not affect a model’s ability to generalize to unseen data (i.e., benign overfitting [54]), but it can lead to inconsistent explanations [55–57], making it difficult to distinguish biologically relevant patterns from spurious importance scores caused by non-biological fluctuations [17, 58]. By averaging attribution maps across an ensemble of models, some of these fluctuations may be reduced, which can lead to more robust explanations [27, 28]. We hypothesized that DEGU-distilled student models, which better approximate the ensemble function, would produce more interpretable and robust attribution maps with stronger motif signals compared to models with standard training.

To test this hypothesis, we generated attribution maps using DeepSHAP [50] and Saliency Maps [52] for models trained with DEGU and with standard training. A visual comparison revealed that the attribution maps from the DEGU-distilled student models displayed more identifiable transcription factor motifs, such as GATA and AP-1 motifs, compared to the models with standard training (Fig. 4a).

**Figure 4.**
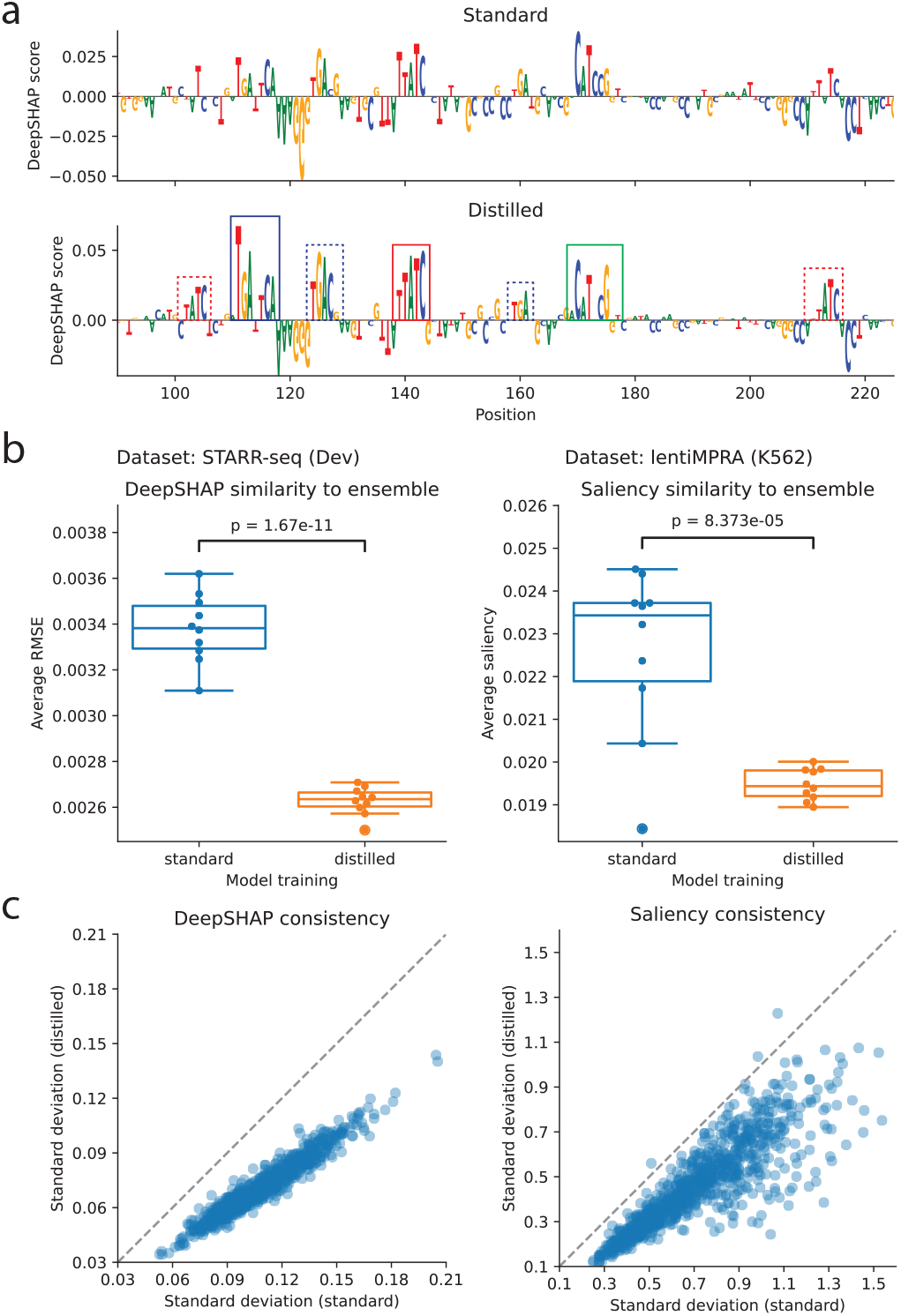
Attribution analysis performance comparison. (**a**) Attribution map for the Dev activity output head of an individual DeepSTARR model with standard training (top) and a DEGU-distilled DeepSTARR model (bottom) for an exemplary test sequence. Annotated boxes indicate binding sites for AP-1 (blue), GATA (red), and ETS/Twist (green), with solid lines indicating a strong match and dashed lines indicating a weak match. (**b**) Average root-mean-squared error (RMSE) between attribution maps generated by individual models with standard training (blue) and DEGU distillation (orange) compared to the average attribution map across the teacher ensemble for DeepSTARR (left) and ResidualBind (right). RMSE is calculated for *n* = 1000 high-activity test sequences. P-values indicate independent two-sided t-tests for average RMSE and paired two-sided t-tests for standard deviation. Box plots represent *n* = 10 models trained with different random initializations, with the boxes representing the first and third quartiles, the central line indicating the median, and whiskers denoting the data range. (**c**) Scatter plots comparing the standard deviation of attribution scores across individual models trained with DEGU distillation (*n* = 10) and standard training (*n* = 10) for DeepSTARR (left) and ResidualBind (right). (**b**,**c**) Attribution scores were calculated with DeepSHAP for the activity output heads of DeepSTARR and with saliency scores for the activity output head of ResidualBind models.

Assuming that ensemble-averaged attribution maps best reflect the underlying biology, we compared the Euclidean distance between attribution maps generated by averaging across the ensemble and those from individual models trained with either standard training or DEGU distillation. We found that the attribution maps produced by distilled models were more closely aligned with the ensemble-averaged attribution maps than those generated by models with standard training across all prediction tasks evaluated (Fig. 4b, Supplementary Fig. 7, Supplementary Fig. 1b).

Additionally, attribution maps generated by different distilled student models were significantly more consistent across random initializations compared to models with standard training (Fig. 4c). This consistency was observed across different models, datasets, and attribution methods (Supplementary Fig. 7, Supplementary Fig. 1b). However, variability across attribution maps from different models can stem from variability in the magnitude of the attribution scores and/or the sequence content (i.e., distinct cis-regulatory mechanisms). To control for variability in attribution score magnitude, we normalized the attribution scores for each sequence. We found that distilled models remained more consistent than the models with standard training (Supplementary Fig. 8), suggesting that the distilled models offer more robust mechanistic insights through their attribution maps.

### DEGU provides calibrated estimates of total uncertainty

A key advantage of deep ensembles is their ability to capture model uncertainty, particularly epistemic uncertainty, which arises from variability in predictions across the ensemble. To evaluate whether DEGU preserves this property, we compared the teacher ensemble’s prediction variability (measured as the standard deviation across predictions) with the epistemic uncertainty estimated by DEGU’s distilled student models. We found that the student models’ predicted uncertainties were strongly correlated with the ensemble’s observed variation, suggesting that DEGU distillation successfully captures epistemic uncertainty (Fig. 5a, Supplementary Fig. 9a, Supplementary Fig. 10). Repeating this analysis with log-variance as the uncertainty measure yielded consistent results (Supplementary Fig. 11).

**Figure 5.**
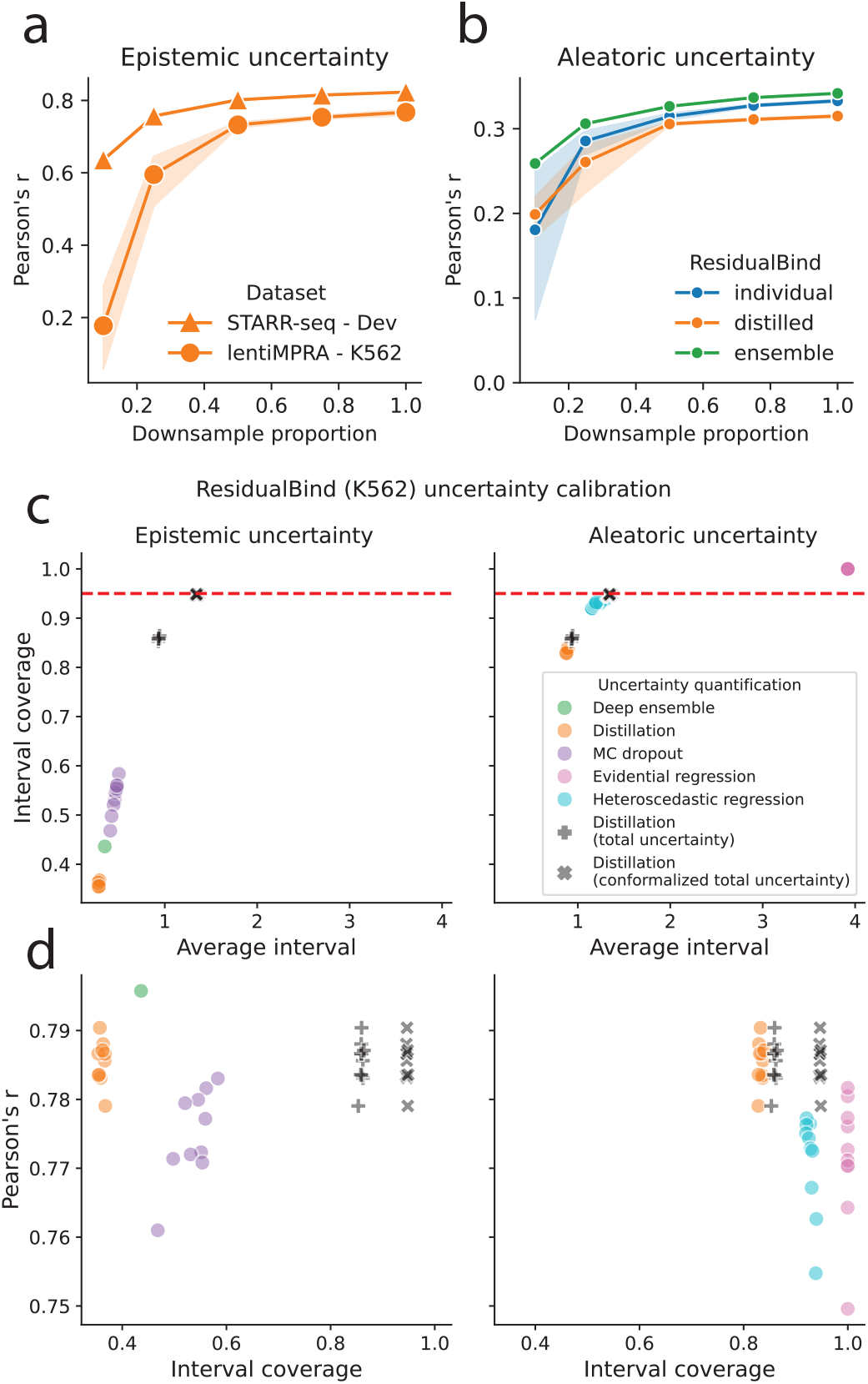
Performance comparison of uncertainty estimates. (**a**,**b**) Predictive performance for (**a**) epistemic and (**b**) aleatoric uncertainty output heads of models trained with standard training (blue), DEGU-distillation (orange), and the teacher ensemble (green), trained on subsets of randomly downsampled training data. Markers represent the average Pearson’s *r* across *n* = 10 models with different random initializations and shaded region indicates 95% confidence interval. Results are shown for the Dev epistemic uncertainty output head of DEGU-distilled DeepSTARR and the uncertainty output heads of ResidualBind models trained on K562 lentiMPRA data. (**c**,**d**) Scatter plots of (**c**) prediction interval coverage probability versus average interval size, and (**d**) predictive accuracy versus interval coverage probability for different methods for quantifying epistemic uncertainty (left) and aleatoric uncertainty (right). Uncertainty quantification methods are based on ResidualBind model trained on K562 lentiMPRA data. Red dashed line indicates calibration with a 95% interval coverage probability. Each uncertainty quantification method is represented by *n* = 10 dots, each indicating a model with different initializations, with the exception of deep ensemble (*n* = 1).

**Figure 6.**
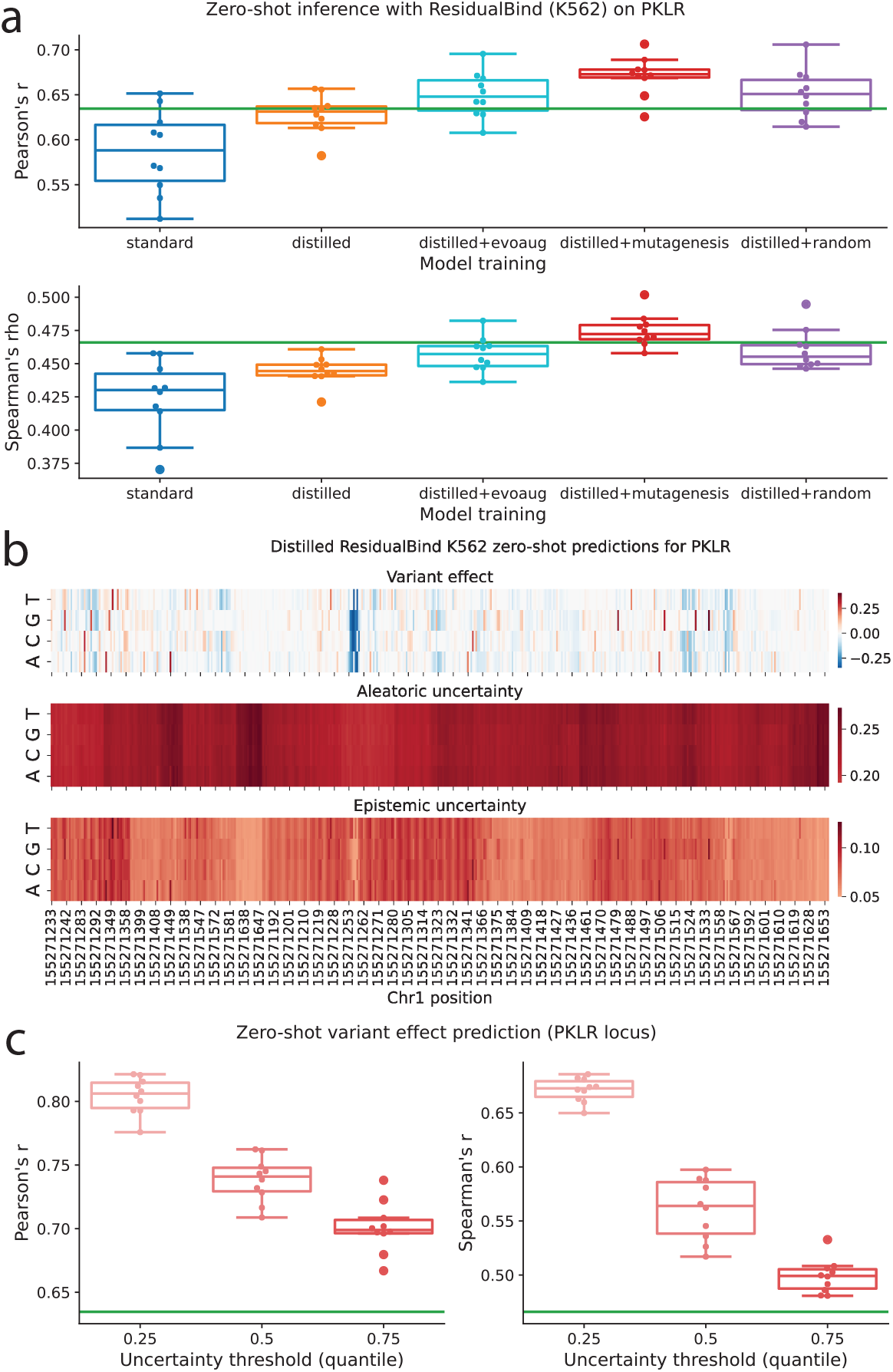
Single-nucleotide variant effect generalization for *PKLR* locus CAGI variants. (**a**) Boxplots of zero-shot variant effect predictive performance for models with standard training, DEGU-distillation, and DEGU-distillation with dynamic augmentations. Predictive performance was evaluated using Pearson’s *r* (top) and Spearman’s *rho* (bottom). (**b**) Heat map showing predicted effect size (top), aleatoric uncertainty (middle), and epistemic uncertainty (bottom) given by a representative distilled ResidualBind model trained on K562 lentiMPRA data. (**c**) Zero-shot variant effect prediction for DEGU-distilled ResidualBind models trained with random mutagenesis augmentations. Predictive performance was evaluated using Pearson’s *r* (left) and Spearman’s *ρ* (right) across *n* = 10 independently initialized models. For each model, variants were stratified by predicted uncertainty, and performance was assessed on subsets of variants below the 25th, 50th, and 75th percentiles of uncertainty. The green horizontal line indicates ensemble teacher performance when evaluated across all variants.

However, relying solely on epistemic uncertainty can lead to overconfident and potentially incorrect predictions [26], as it does not capture the full spectrum of predictive uncertainty. DEGU addresses this limitation by training models to also predict aleatoric uncertainty, which is estimated from the variability (measured as the standard deviation) across experimental replicates (when sufficient replicates are available). We demonstrated this approach using the lentiMPRA [23] and ATAC-seq profile datasets, both of which had three experimental replicates. We assessed the performance of our models’ aleatoric uncertainty predictions using Pearson’s r and found that predicting aleatoric uncertainty from sequence data remains challenging (Fig. 5b, Supplementary Figs. 9b and 1c), which is expected given that aleatoric uncertainty represents irreducible noise. While accurately predicting aleatoric uncertainty is inherently challenging, ensuring that the predicted uncertainties are well-calibrated is more important.

To evaluate the reliability of DEGU’s uncertainty estimates, we performed an uncertainty calibration analysis using the prediction interval coverage probability. This metric measures how often the true target value for a given sequence (i.e., experimental values for regulatory activity) falls within the confidence interval estimated for the corresponding predicted values for activity and uncertainty. Ideally, a well-calibrated model would achieve a coverage probability equal to the confidence interval percentage (95% in this analysis; see Methods), while minimizing the size of the interval.

We benchmarked the calibration of uncertainty estimates from models trained with DEGU against various other uncertainty quantification strategies, including those that capture epistemic uncertainty, such as Monte Carlo Dropout (MCDropout) [59] and deep ensembles, as well as methods for estimating aleatoric uncertainty, such as deep evidential regression [60] and heteroscedastic regression [61, 62]. Our results showed that methods accounting for aleatoric uncertainty, including DEGU’s aleatoric head and heteroscedastic regression, generally achieved better calibration with smaller average interval sizes. In contrast, methods relying solely on epistemic uncertainty (e.g., DEGU’s epistemic head or MCDropout) exhibited severe under-calibration (Fig. 5c, Supplementary Fig. 9c).

We also observed a trade-off between predictive accuracy and uncertainty calibration across different methods (Fig. 5d, Supplementary Fig. 9d). Models trained with heteroscedastic regression were well-calibrated but had slightly lower predictive accuracy compared to distilled models. Conversely, distilled models achieved high predictive performance but with slightly worse uncertainty calibration with respect to total uncertainty (Fig. 4c-d, Supplementary Fig. 9c-d).

Since ResidualBind models trained with heteroscedastic loss produced better-calibrated estimates of aleatoric uncertainty compared to models that predicted aleatoric uncertainty based on replicate-level variation but yielded lower predictive performance for sequence activity, we next investigated whether applying DEGU distillation to models trained with a heteroscedastic regression loss could close this performance gap. However, this approach did not fully bridge the gap between heteroscedastic regression and DEGU (Supplementary Fig. 12).

To improve calibration, we applied conformal prediction [63–65] to the uncertainty estimates, which ensures guaranteed coverage under minimal assumptions of data exchangeability. This resulted in nearly perfect calibration for all uncertainty quantification methods evaluated (Fig. 5c, Supplementary Fig. 13), giving the distilled models the best overall performance, with high predictive accuracy and well-calibrated uncertainty estimates.

### Uncertainty-aware zero-shot variant effect generalization

Uncertainty quantification offers a valuable tool for informed decision making, particularly in scenarios where the reliability of model predictions is unclear, such as single-nucleotide variant effect prediction. Previously, we demonstrated that data augmentations improved function approximation to the ensemble (Supplementary Fig. 6), suggesting potential for better generalization. To directly test this, we trained distilled ResidualBind models with dynamic augmentations on lentiMPRA data and evaluated their ability to predict single-nucleotide variant effects in matched cell types using experimental measurements from massively parallel reporter assays (MPRAs) [66, 67].

As expected, variant effect predictions from the distilled models generally showed stronger correlation with experimentally measured variant effects compared to models with standard training, with additional performance gains when distilled models were trained with data augmentations (Fig. a, Supplementary Fig. 14). Visual analysis indicated that aleatoric uncertainty predictions were greater in magnitude and more consistent across nucleotides at a given position, whereas epistemic uncertainty predictions exhibited greater variability across both nucleotides and positions (Fig. b). Further analysis revealed that aleatoric uncertainty was higher for variant effects close to zero, while epistemic uncertainty increased as activity levels moved further from zero (Supplementary Fig. 15).

To investigate the relationship between uncertainty and predictive accuracy, we stratified model performance based on a total uncertainty threshold, classifying predictions below this threshold as confident (see Methods). We focused on distilled ResidualBind models trained with random mutagenesis augmentations, as they yielded the best overall performance (Supplementary Fig. 14). This uncertainty-based stratification revealed that the most confident predictions were associated with higher performance (Fig. c), although the trends varied across different loci (Supplementary Fig. 16). These findings highlight the potential of uncertainty quantification to enhance the reliability and interpretability of variant effect predictions, enabling more nuanced analysis and decision making in genomics research.

## Discussion

DEGU presents a simple and effective approach for harnessing the benefits of deep ensembles with just a single model, providing robust predictions alongside reliable uncertainty estimates. Our results demonstrate that DEGU-distilled models generally outperform models with standard training and provide more reliable post hoc explanations of cis-regulatory mechanisms through attribution analysis. This makes DEGU particularly well-suited for large-scale inference tasks, such as genome-wide variant effect prediction and generating post hoc explanations across a large number of regulatory sequences.

A major strength of DEGU lies in its dual uncertainty estimation, addressing a key limitation of current genomic deep learning models. By simultaneously estimating both epistemic and aleatoric uncertainty, DEGU offers a comprehensive assessment of prediction reliability – an important factor in enhancing confidence in model predictions.

DEGU excels in estimating epistemic uncertainty, which was straightforward to train using prediction variability across the teacher ensemble. In contrast, estimating aleatoric uncertainty posed greater challenges due to inherent randomness of data noise. To approximate this, we created a prediction task based on experimental replicates, yielding aleatoric estimates with well-calibrated prediction intervals that surpassed those based on epistemic uncertainty. We also evaluated heteroscedastic regression as an alternative approach for aleatoric uncertainty, which offers a simpler method that avoids extra data processing steps and an additional prediction task. However, heteroscedastic regression yielded slightly reduced predictive performance on functional activities and proved challenging to optimize due to an unstable loss function. Overall, aleatoric uncertainty estimates based on replicate variability combined with epistemic uncertainty estimates achieved the best balance between calibration and predictive accuracy. Nevertheless, heteroscedastic regression loss can serve as an alternative approach when there are insufficient replicates.

DEGU’s ability to approximate the teacher ensemble’s function suggests that distilled models generalize better under covariate shifts. Moreover, training with dynamic data augmentations led to improved approximation of the ensemble. This proved to be an effective approach to improve zero-shot predictions of single-nucleotide variant effects. Since most downstream applications of genomic DNNs (i.e. single-nucleotide variant effect prediction, counterfactual in silico perturbation experiments, and synthetic DNA sequence design) involve varying degrees of covariate shifts, DEGU-distilled models are well suited to provide robust predictions with uncertainty estimates that reflect its confidence.

An important limitation of our study is that we evaluated DEGU only on single-nucleotide variant effect prediction tasks; however, prior work has highlighted that population-level sparse mutations remain especially challenging for current models [18, 19]. Extending DEGU to effectively capture uncertainty and improve prediction accuracy in this setting is a promising avenue for future research, particularly given the relevance of personalized genetic variation to disease risk and regulatory function.

Our study focused solely on ensembles of models with the same architecture. There is also great potential in applying DEGU to more compact student architectures, which could improve the efficiency of deploying large-scale DNNs such as Enformer [1] and Borzoi [68], both of which currently have high computational costs. Making these models more computationally efficient would reduce the need for extensive GPU resources, thereby democratizing access to state-of-the-art genomic prediction tools. Further improvements could be achieved by increasing the diversity of models within the ensemble and experimenting with different weighting methods for ensemble members.

In this study, DEGU primarily focused on approximating the ensemble’s function, leaving opportunities for further improvement by incorporating a mixed knowledge distillation loss function that balances the use of real training data with the divergence between the ensemble and student models [37]. A related approach to DEGU is self-distillation [69], where the distillation process is carried out sequentially in an online manner. While self-distillation can offer benefits similar to ensemble distillation [70], it inherently struggles to effectively capture epistemic uncertainty, as it relies on a single model’s training path rather than the diverse perspectives provided by an ensemble. In contrast, DEGU explicitly models epistemic uncertainty (and aleatoric uncertainty) as part of its distillation process. In the future, we plan to gain deeper insights into the sequence features that contribute to uncertainty in genomic predictions through a comprehensive attribution analysis on the uncertainty head.

DEGU’s ability to provide insights into the confidence of each prediction represents an important step toward making deep learning models more reliable and trustworthy. While our demonstration of DEGU focused on deep learning models for regulatory genomics, the process of ensemble distribution distillation is general and can be extended to other models in other domains of biology. As the field continues to evolve, uncertainty-aware models like DEGU will likely become essential for guiding research decisions and clinical applications, highlighting the importance of further refining and expanding these techniques.

## Methods

### Datasets

#### Fly enhancer activity with STARR-seq

We obtained STARR-seq data for developmental (Dev) and housekeeping (Hk) promoters in *D. melanogaster* S2 cells from de Almeida et al.[40] Each sequence is 249 base pairs (bp) long. Enhancer activity for both housekeeping and developmental classes was predicted simultaneously as a multi-task regression. The data was split into train, test, and validation sets containing 402296, 41186, and 40570 samples, respectively.

#### Human regulatory sequences with lentiMPRA

We used lentiMPRA data for K562 and HepG2 cell lines from Agarwal et al.[23]. Each 230 bp cis-regulatory sequence was associated with a scalar activity measurement for three biological replicates. The mean and standard deviation across the replicates was used as target values for regulatory sequence activity and aleatoric uncertainty, respectively. For each cell type, we performed two types of regressions: 1) a single-task regression for regulatory activity only, and 2) a multi-task regression for both regulatory activity and aleatoric uncertainty. We generated a different dataset for each regression task. For the single-task regression, we removed any samples for which an activity measurements was provided without corresponding sequence data was not available. For the multi-task regression, we also removed samples for which experimental data from at least two replicates was not available, due to the inability to calculate aleatoric uncertainty. For each dataset, we randomly split the training, validation, and test sets according to the fractions 0.8, 0.1, and 0.1, respectively, ensuring that any forward and reverse complement sequence pairs would be assigned to the same set to avoid data leakage. The HepG2 data for single-task regression (activity only) was split into train, test, and validation sets containing 111901, 13988, and 13988 samples, respectively. The HepG2 data for the multi-task regression (activity and aleatoric uncertainty) was split into train, test, and validation sets containing 111518, 13939, and 13942 samples, respectively. The K562 data for single-task regression (activity only) was split into train, test, and validation sets containing 181002, 22626, and 22626 samples, respectively. The K562 data for multi-task regression (activity and aleatoric uncertainty) was split into train, test, and validation sets containing 180564, 22571, and 22570 samples, respectively.

#### Profile-based chromatin accessibility with GOPHER

We acquired hg38-aligned ATAC-seq bigWig files for A549 cells (ENCSR032RGS) from ENCODE [71]. We processed the 3 replicate fold change over control bigwig files into 2 bigWig files: average read coverage across replicates and 2) standard deviation of read coverage across replicates. Wiggletools [72] and UCSC’s bedGraphToBigWig [73] were used to wrangle the data into bigWig file formats. Following a previously published data processing procedure [8], we divided each chromosome into equal, non-overlapping 3072 bp bins. We one-hot encoded each sequence with matched base-resolution coverage tracks from the average and standard deviation bigWig files. We split the dataset into a test set comprising chromosome 8, a validation set comprising chromosome 9, and a training set encompassing the remaining chromosomes, with the exclusion of chromosome Y and contigs. Performance was assessed as the Pearson correlation across the whole chromosome as outlined in Ref. [8].

#### Single-nucleotide variant effect with CAGI5

The CAGI5 challenge dataset [66, 67], which consists of experimentally measured saturation mutagenesis of a 230 bp regulatory element via a MPRA, was used to evaluate the performance of the ResidualBind models on zero-shot single-nucleotide variant effect generalization. We considered only experiments in HepG2 (*LDLR, F9, SORT1*) and K562 (*PKLR*). We extracted 230 bp sequences from the reference genome (hg19) centered on each single-nucleotide variant in the CAGI data. We calculated the predicted effect of each allele as: ŷ_*alt*_ ŷ_*ref*_, where ŷ_*alt*_ is the model’s activity prediction for the alternate allele and ŷ_*ref*_ is the model’s activity prediction for the corresponding reference allele. Performance was evaluated as the Pearson correlation between the predicted effect and the experimentally measured effect.

### Models

#### DeepSTARR

We implemented DeepSTARR [40] as described in Ref [40], according to:

1. 1D convolution (256 kernels, size 7, batch normalization, ReLU activation) 1D max-pooling (size 2)
2. 1D convolution (60 kernels, size 3, batch normalization, ReLU activation) 1D max-pooling (size 2)
3. 1D convolution (60 kernels, size 5, batch normalization, ReLU activation) 1D max-pooling (size 2)
4. 1D convolution (120 kernels, size 3, batch normalization, ReLU activation) 1D max-pooling (size 2)
5. flatten
6. linear (256 units, batch normalization, ReLU activation) dropout(0.4)
7. linear (256 units, batch normalization, ReLU activation) dropout(0.4)
8. output (2 units, linear)

The 2 units in the output layer represent the Dev and Hk enhancer activities. For distilled models which predict both activity and epistemic uncertainty, the output layer is increased to 4 units, representing Dev activity, Hk activity, Dev epistemic uncertainty, and Hk epistemic uncertainty.

#### ResidualBind for lentiMPRA

We used a custom ResidualBind model[6, 42], a CNN with dilated residual blocks [74, 75], to model lentiMPRA data. The ResidualBind architecture is as follows:

1. 1D convolution (196 kernels, size 19, batch normalization, SiLU activation) dropout (0.2)
2. Dilated residual block (5 dilations) 1D convolution (196 kernels, size 3, batch normalization, ReLU activation) dropout (0.1) 1D convolution (196 kernels, size 3, dilation rate 1, batch normalization, ReLU activation) dropout (0.1) 1D convolution (196 kernels, size 3, dilation rate 2, batch normalization, ReLU activation) dropout (0.1) 1D convolution (196 kernels, size 3, dilation rate 4, batch normalization, ReLU activation) dropout (0.1) 1D convolution (196 kernels, size 3, dilation rate 8, batch normalization, ReLU activation) dropout (0.1) 1D convolution (196 kernels, size 3, dilation rate 16, batch normalization, ReLU activation) dropout (0.1) skip connection to input SiLU activation dropout (0.2) 1D max-pooling (size 5)
3. 1D convolution (256 kernels, size 7, batch normalization, SiLU activation) dropout (0.2) 1D max-pooling (size 5)
4. linear (256 units, batch normalization, SiLU activation) dropout(0.5)
5. 1D global average pooling
6. flatten
7. linear (256 units, batch normalization, SiLU activation) dropout(0.5)
8. output (1 unit, linear)

For ResidualBind models trained on both the replicate average and standard deviation, the output layer is increased to 2 units representing activity and aleatoric uncertainty, respectively. For distilled ResidualBind models, the output layer is increased to 3 units representing activity, aleatoric uncertainty, and epistemic uncertainty.

#### CNN-task-base

A base-resolution CNN from Ref. [8] was used to fit the ATAC-seq profile data. Briefly, CNN-task-base is composed of 3 convolutional blocks, which consist of a 1D convolution, batch normalization, activation, max pooling and dropout, followed by 2 fully-connected blocks, which includes a dense layer, batch normalization, activation, and dropout. The first fully connected block scales down the size of the representation, serving as a bottleneck layer. The second fully-connected block rescales the bottleneck to the target resolution. This is followed by another convolutional block. The representations from the outputs of the convolutional block is then input into task-specific output heads; each head consists of a convolutional block followed by a linear output layer with softplus activations. The activation of all hidden layers are ReLU.

#### DREAM-RNN

We used a custom DREAM-RNN model, a hybrid architecture combining convolutional and recurrent layers [43]. The DREAM-RNN architecture is as follows:

1. Parallel 1D convolution block 1D convolution (256 kernels, size 9, ReLU activation) dropout (0.2) 1D convolution (256 kernels, size 15, ReLU activation) dropout (0.2) concatenate outputs along channel dimension
2. Bidirectional LSTM (320 hidden units per direction, concatenated output of 640 dimensions) dropout (0.2)
3. Parallel 1D convolution block 1D convolution (256 kernels, size 9, ReLU activation) dropout (0.2) 1D convolution (256 kernels, size 15, ReLU activation) dropout (0.2) concatenate outputs along channel dimension
4. point-wise 1D convolution (256 kernels, size 1, ReLU activation) dropout (0.2)
5. 1D global average pooling
6. flatten
7. linear (units depending on output task) SoftMax (for classification) or linear activation (for regression)

For multi-task DREAM-RNN models, the output layer is expanded as needed (e.g., 2 units for activity and aleatoric uncertainty; 3 units for activity, aleatoric uncertainty, and epistemic uncertainty).

### Training models

#### Standard training of DeepSTARR, ResidualBind and DREAM-RNN

We uniformly trained each model by minimizing the mean-squared error loss function with mini-batch stochastic gradient descent (100 sequences) for 100 epochs with Adam updates using default parameters [76]. The learning rate was initialized to 0.001 and was decayed by a factor of 0.1 when the validation loss did not improve for 5 epochs. All reported performance metrics are drawn from the test set using the model parameters from the epoch which yielded the lowest loss on the validation set. For each model, we trained 10 different individual models with different random initializations.

#### Standard training of CNN-task-base

CNN-task-base models were trained using a Poisson loss and Adam with default parameters and a minibatch size of 100. The learning rate was initialized to 0.001 and was decayed by a factor of 0.3 when the validation loss did not improve for 5 epochs. All reported performance metrics are drawn from the test set using the model parameters from the epoch which yielded the lowest loss on the validation set. During training, random shift and stochastic reverse-complement data augmentations were used [8]. Random shift is a data augmentation that randomly translates the input sequence (and corresponding targets) online during training. For each mini-batch, a random sub-sequence of 2048 bp and its corresponding target profile was selected separately for each sequence. Reverse-complement data augmentation is also employed online during training. During each mini-batch, half of training sequences were randomly selected and replaced by their reverse-complement sequence. For those sequences that were selected, the training target was correspondingly replaced by the reverse of original coverage distribution. For each model, we trained 10 different individual models with different random initializations.

#### Training DeepSTARR with EvoAug

Models trained with EvoAug-TF [45] use the following augmentation settings:

- random deletions with a size range of 0-20bp (applied per batch)
- random translocation with a size range of 0-20bp (applied per batch)
- random Gaussian noise with *µ* = 0 and *σ* = 0.2 added to each variant in the input sequence (applied per sequence)
- random mutation of 5% of nucleotides in sequence (applied per sequence)

For each minibatch during training, one of the augmentations is randomly selected from the list of possible augmentations described above and applied to every sequence in the minibatch. Both teacher and student models were trained with the same optimizer, learning rate decay, and early stopping hyperparameters described for standard training.

### DEGU: distilling knowledge of ensembles to uncertainty-aware genomic DNNs Ensemble Training

For each prediction task, we trained an ensemble of *M* models 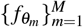 with identical architectures but different random initializations *θ*_*m*_. For an input sequence *x*_*i*_ (with *i* = 1, …, *N*), the ensemble yields predictions 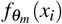. We summarize the ensemble by its mean and standard deviation:

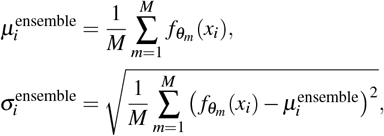

where 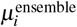 is the ensemble mean prediction and *σ* ^ensemble^ captures ensemble variability (used as labels for epistemic uncertainty).

#### Distilled Model Architecture

The distilled model *g*_*φ*_ consists of a shared trunk *T* (·; *φ*_trunk_) that maps each sequence *x*_*i*_ to a latent representation:

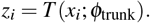

From this shared representation, we apply two output heads for epistemic prediction:

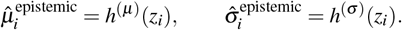

Thus, in the basic setting, the distilled model outputs

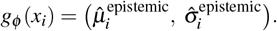

Each head has its own parameters, distinct from the trunk and from one another. This formulation is general and does not assume any particular functional form for the heads. In this paper, we instantiate all output heads as linear layers.

#### Loss Function

We train the distilled model to match the ensemble-derived labels:

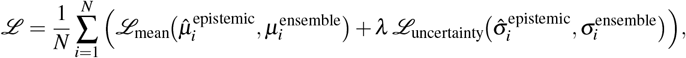

where both losses are mean squared error terms, and *λ* = 1 in this study.

#### Aleatoric Uncertainty (Optional)

When experimental replicates are available (*R*_*i*_ ≥ 3), we can also estimate aleatoric uncertainty directly from the variability across replicates. Let 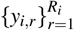denote replicate measurements for input *x*_*i*_. The replicate variance provides a label for aleatoric uncertainty:

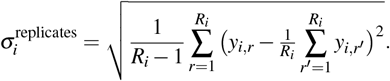

To capture this variability, we add an aleatoric uncertainty head:

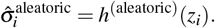

The full model then outputs

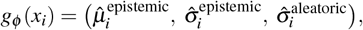

and the loss is augmented as

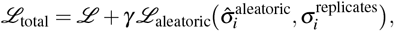

with *γ* = 1 in this study.

#### Model Architecture

While the distilled models can be comprised of any architecture, in this study, the distilled models share the same architecture as the original ensemble models, with the exception of the final layer. If the original models had *N*_out_ output heads, the distilled models will have 2*N*_out_ heads to account for both the mean and epistemic uncertainty predictions. In cases where aleatoric uncertainty is also modeled, the distilled models will have 3*N*_out_ output heads.

#### Distilling DeepSTARR and DREAM-RNN

We first trained an ensemble of 10 models with different random initializations on STARR-seq data. We then used each individual model in the ensemble to make predictions on the training sequences and calculated the average and standard deviation across the 10 models. These values were used as target values for activity and epistemic uncertainty, respectively. Then, we trained 10 distilled models with different random initializations following the same procedure as standard training using the new target labels generated by the teacher ensemble.

#### Distilling ResidualBind and DREAM-RNN

For each cell type, we also trained an ensemble of 10 models with different random initializations on both the mean and standard deviation of experimental activity values across the biological replicates in the lentiMPRA data, with the latter value representing aleatoric uncertainty. The averages of the activity and aleatoric uncertainty predictions from this ensemble were used as new target values for training the distilled models. Moreover, the standard deviation of the activity predictions were also used to generate labels for the epistemic uncertainty. Then, we trained 10 distilled models with different random initializations following the same procedure as standard training using the new target labels generated by the teacher ensemble.

#### Distilling CNN-task-base

We first trained an ensemble of 10 CNN-task-base models with different random initializations on profile-based ATAC-seq data. These models were trained to predict the mean and standard deviation of the profiles across the three biological replicates. We then used the ensemble of models to generate new bigWig tracks based on the mean and standard deviation of the profiles across the 10 models, using wiggletools. We also included the mean of the aleatoric uncertainty head, resulting in 3 bigWig tracks all generated by the predictions of the ensemble. Each bigWig was processed following the same procedure as the original ATAC-seq profiles. Distilled models used the same architecture and training procedure as the standard CNN-task-base with the exception of using ensemble-generated labels and the addition of the epistemic profile prediction task, resulting in 3 prediction tasks, including the mean profile, the aleatoric uncertainty profile, and the epistemic uncertainty profile. Then, we trained 10 distilled models with different random initializations following the same procedure as standard training using the new target labels generated by the teacher ensemble.

#### Distilling models with dynamic augmentations

During distillation, we generated data augmentations following three different augmentation schemes:

1. EvoAug-TF[45], with *n* augmentations selected from the same augmentation list described above where *n* is randomly selected from 0 to 2.
2. Random mutagenesis using the EvoAug-TF implementation with a mutation fraction of 5%
3. Random shuffling

The augmentation is applied to each minibatch during training, replacing the original training sequences. The ensemble of teacher models is used to make predictions on the augmented sequences, and the average and standard deviation of these predictions are used as target values for activity and uncertainty, respectively. The original validation and test sequences were used for early stopping and evaluations. Student models were trained with the same optimizer, learning rate decay, and early stopping hyperparameters described for standard training.

### OOD generalization analysis

We generated out-of-distribution (OOD) sequences using the following sampling methods:

1. Small distribution shift: random mutagenesis with a mutation fraction of 5% generated with EvoAug[44].
2. Moderate distribution shift: evolution-inspired mutagenesis generated with 2 augmentations selected from the same augmentation list described above using EvoAug[44]
3. Large distribution shift: random shuffling of the test sequences.

Activity labels for these OOD sequences were obtained by averaging the predictions from an *in silico* oracle comprised of an ensemble of DeepSTARR models trained with EvoAug (with the same hyperparameters as stated above).

### Attribution analysis

Saliency Maps [52] and DeepSHAP [50] scores were employed for attribution analysis to elucidate the input nucleotides most influential model predictions. For each sequence activity output head of each model, we generated attribution maps for 1000 sequences from the test set associated with the largest target values. Each method yielded a 4 × *L* map where *L* is the length of the input sequence. For DeepSHAP, background sequences were comprised of 100 randomly selected sequences from the test set. Gradient correction was applied to all attribution maps by subtracting the average attribution score across all channels (nucleotides) at each position [28]. For profile-based models, we transformed the predictions to a scalar through a global average along the length dimension.

#### Calculating similarity of attribution scores to ensemble average

For each individual model, we calculated the root mean squared error (RMSE) of the attribution maps for the 1000 sequences evaluated between the individual models and the average attribution maps of the teacher ensemble. Individual models refer to those trained with either DEGU distillation or standard training from random initializations. The teacher ensemble was calculated by averaging the attribution maps across the 10 individual models with standard training.

#### Calculating variability of attribution scores across different initializations

For each of the 1000 sequences evaluated, we calculated the variance of their attribution scores for each nucleotide and position in each sequence across 10 individual models (trained with different random initializations). These per-nucleotide and per-position variances are then summed across the sequence to calculate the total variance, followed by a square root operation to provide a measure of the standard deviation of attribution scores across different initializations.

#### Control experiment with normalized attribution scores

For the control experiment that isolated mechanistic variability, we obtained the per-sequence maximum attribution score magnitude across nucleotides and positions for each of the 1000 sequences evaluated and then divided all attribution scores by this value to obtain an attribution-magnitude normalized set of attribution score.

### Evaluating the size of the teacher ensemble for DeepSTARR

We trained an additional 15 DeepSTARR models using different random initializations using the entire STARR-seq training set for a total of 25 models. We then performed ensemble distribution distillation for subsets of 2, 3, 4, 5, 15, and 20 of these replicates, as well as for the entire set of 25 replicates. We evaluated the predictive accuracy of the activity predictions for the individual models in these ensembles, the ensemble average, and the distilled models derived from the respective teacher ensembles and compared them across different teacher ensemble sizes.

### Uncertainty-aware models

#### Heteroscedastic regression

ResidualBind models trained with heteroscedastic regression utilized a Gaussian negative log-likelihood loss function. The final output layer was modified to a linear layer with 2 output heads representing the mean (*µ*) and log variance (log *σ* ^2^). The use of log variance ensures numerical stability during training and guarantees positive variance predictions. The loss function is defined as:

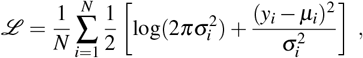

where *µ*_*i*_ and 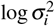 are predicted by the model and *N* is the number of samples in the batch. The model was trained using mini-batch stochastic gradient descent with the same optimizer, learning rate decay, and early stopping settings as used for standard training of the models described above. The variance predicted by heteroscedastic regression represents aleatoric uncertainty.

#### Deep evidential regression

ResidualBind models trained with deep evidential regression[60] were modified so that their output layer considers the mean (*µ*) and log-variance (log *σ* ^2^), which represents an estimate of the aleatoric uncertainty. The loss function is defined as:

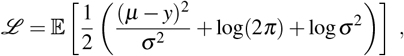

where 𝔼 [·] denotes the expectation (average) over all dimensions and samples in the batch. The terms *µ* and log *σ* ^2^ are predicted by the model and *y* is the true value.

#### MC Dropout

We implemented Monte Carlo (MC) dropout as described by Gal and Ghahramani [59]. This method leverages dropout at inference time to estimate predictive uncertainty. For each input, we performed 100 stochastic forward passes through the model, with dropout remaining active during inference. We then calculated the mean and standard deviation across the predictions for each input. The mean prediction represents the model’s best estimate, while the standard deviation quantifies the epistemic uncertainty associated with that prediction.

### Uncertainty calibration analysis

#### Interval coverage analysis

For each uncertainty quantification method, we calculated a 95% confidence interval using the model’s prediction of sequence activity and uncertainty for the test sequences in model’s corresponding dataset. For uncertainty quantification methods that yielded both aleatoric and epistemic uncertainty estimates, we calculated intervals based on each of the two different uncertainty estimates as well as the total uncertainty calculated as the sum of the variances, according to:

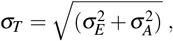

where *σ*_*T*_ is total uncertainty, 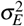 is epistemic uncertainty, and 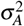 is aleatoric uncertainty. For models where the uncertainty prediction was given as log-variance (i.e. models trained with heteroscedastic regression), the output was accordingly transformed for compatibility with total uncertainty as a measure of standard deviation. The interval coverage probability was calculated as the fraction of cases where the experimental activity value fell within the 95% confidence interval constructed from the predicted activity and uncertainty values. Assuming a Gaussian distribution, the 95% confidence interval was calculated as 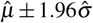, where 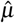 and 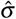 represent the estimates of activity and uncertainty for the method being evaluated.

#### Conformal predictions

Conformal prediction was used to calibrate the predicted uncertainties, according to:

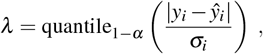

where *y*_*i*_ are the true target values for the calibration sequence *i*, ŷ_*i*_ are the predicted values for the calibration sequence *i*, 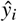 are the uncertainty estimates for the calibration sequence *i*, and *α* is the desired confidence threshold, set to 0.05.

#### Calibration sequences were taken from the validation set

The calibration factor *λ* is then multiplied by the predicted uncertainty estimates for the test sequences.

## Data Availability

Processed data and model weights can be found at DOI:10.5281/zenodo.14145284. Datasets include lentiMPRA, STARR-seq, ATAC-seq profile data, and zero-shot single-nucleotide variant effect data.

## Code Availability

Open-source code to reproduce this study is available on GitHub (https://github.com/zrcjessica/ensemble_distillation).

## Acknowledgements

The authors would like to thank Moon (Masayuki) Nagai for helpful comments on the manuscript. Research reported in this publication was supported in part by the National Human Genome Research Institute of the National Institutes of Health under Award Number R01HG012131 and the National Institute Of General Medical Sciences of the National Institutes of Health under Award Number R01GM149921. This work was performed with assistance from the US National Institutes of Health Grant S10OD028632-01.

## Author contributions

JZ and PKK conceived of the method and designed the experiments. JZ developed code, ran the experiments, and analyzed the results for DeepSTARR and ResidualBind models. KR and ZT performed the base-resolution ATAC-seq profile analysis. TC trained DREAM-RNN models and benchmarked model inference times. JZ and PKK interpreted the results. JZ, PKK, KR, and TC contributed to writing the paper.

---

**Supplementary Table 1.** Inference time comparison across different model architectures and datasets. Results show mean inference times and proportional comparisons between the distilled model and ensemble/teacher models per batch (1024 sequences) averaged over 20 timed trials following 10 warmup runs. All experiments conducted on NVIDIA A3000 GPU with CUDA optimization and cache stabilization. Ratios compare distilled model inference times versus ensemble and standard architectures.

| **Dataset** | **Standard**<br>**Time (s)** | **Ensemble**<br>**Time (s)** | **Distilled**<br>**Time (s)** | **Distilled /**<br>**Ensemble** | **Distilled /**<br>**Standard** |
| --- | --- | --- | --- | --- | --- |
| DeepSTARR | 0.145 | 1.469 | 0.147 | 0.100 | 1.01 |
| lentiMPRA_K562 | 1.151 | 11.480 | 1.150 | 0.100 | 1.00 |
| lentiMPRA_HepG2 | 1.153 | 11.459 | 1.146 | 0.100 | 0.99 |

**Supplementary Figure 1.**
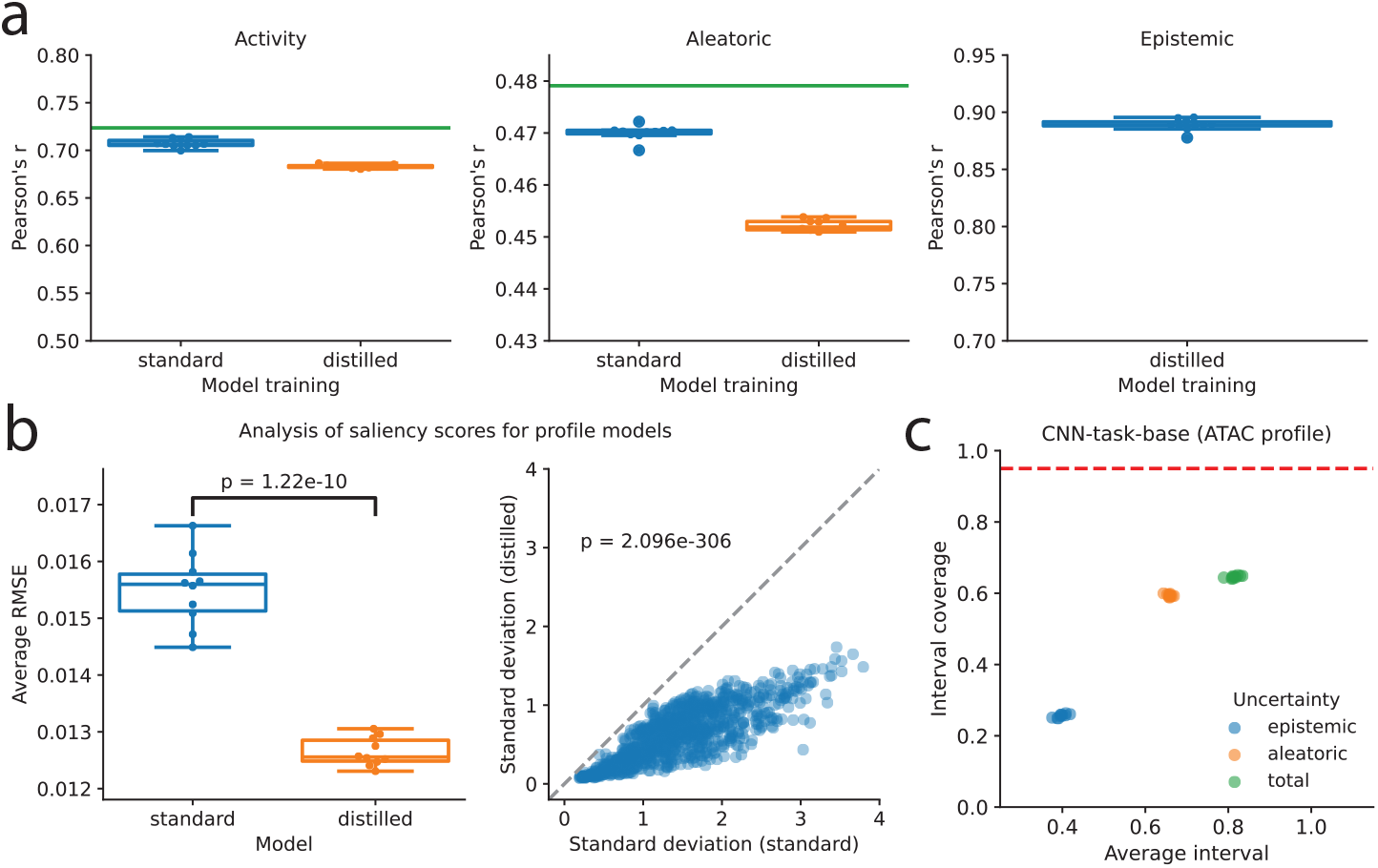
Performance on ATAC-seq profiles. (**a**) Boxplots of prediction performance of base-resolution CNN model trained on ATAC-seq profiles with standard training (blue) and DEGU distillation (orange) for the activity head (left), aleatoric head (middle) and epistemic head (right). Green horizontal line indicates the teacher ensemble performance. Boxplots represent *n* = 10 models trained with different random initializations, with the boxes indicating the first and third quartiles, the central line indicates the median, and whiskers denote the data range. (**b**) (left) Boxplot of average root mean squared error (RMSE) between attribution maps generated by individual models trained with standard training (blue) and DEGU distillation (orange), compared to the average attribution map across the teacher ensemble, and (right) scatter plot comparing the standard deviation of attribution scores across individual models trained with standard training (*n* = 10) and DEGU distillation (*n* = 10). Attribution scores were calculated with Saliency Maps for the activity output. RMSE is calculated for 1,000 high-activity test sequences. P-values indicate independent two-sided t-tests for average RMSE and paired two-sided t-tests for standard deviation. (**a**,**b**)Boxplots represent *n* = 10 models trained with different random initializations, with the boxes representing the first and third quartiles, the central line indicating the median, and whiskers denoting the data range. (**c**) Scatter plot of prediction interval coverage probability versus average interval size for epistemic uncertainty (blue) and aleatoric uncertainty (orange) and total uncertainty (green). Red dashed line indicates calibration with a 95% interval coverage probability. Each uncertainty quantification method is represented by *n* = 10 dots, indicating a model with different initializations, except for deep ensemble (*n* = 1).

**Supplementary Figure 2.**
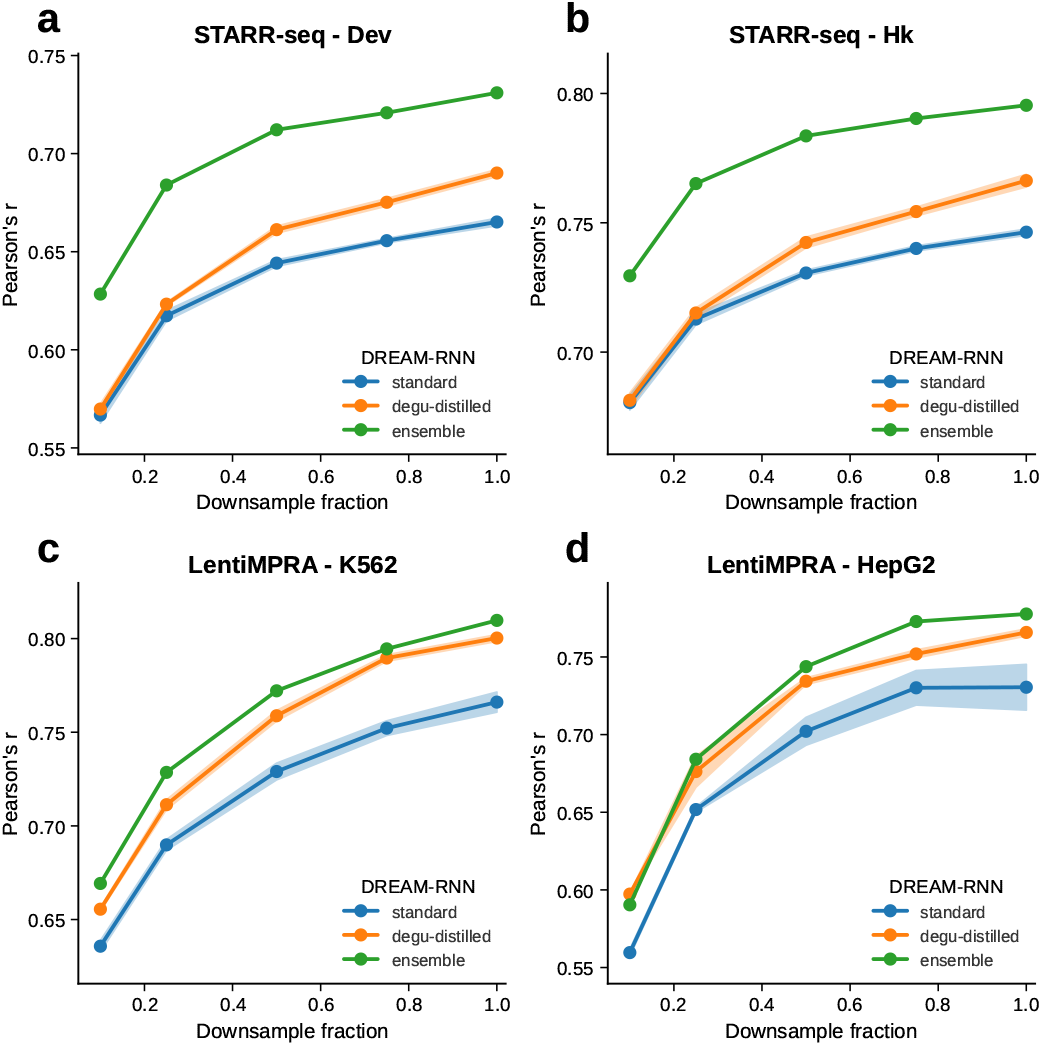
Comparison of model performance with downsampled training data with DREAM-RNN. (**a**,**b**) Predictive performance of DREAM-RNN models trained on different subsets of randomly downsampled STARR-seq data for (**a**) developmental (Dev) and (**b**) housekeeping (Hk) promoters. (**c**,**d**) Predictive performance of DREAM-RNN models trained on subsets of randomly downsampled lentiMPRA data for(**c**) K562 and (**d**) HepG2 cell lines. (**a**-**d**) Plots include models with standard training (blue; *n* = 10), DEGU-distillation (orange; *n* = 5); and the ensemble average of the teacher models with standard training (green). Shaded regions indicate 95% confidence intervals.

**Supplementary Figure 3.**
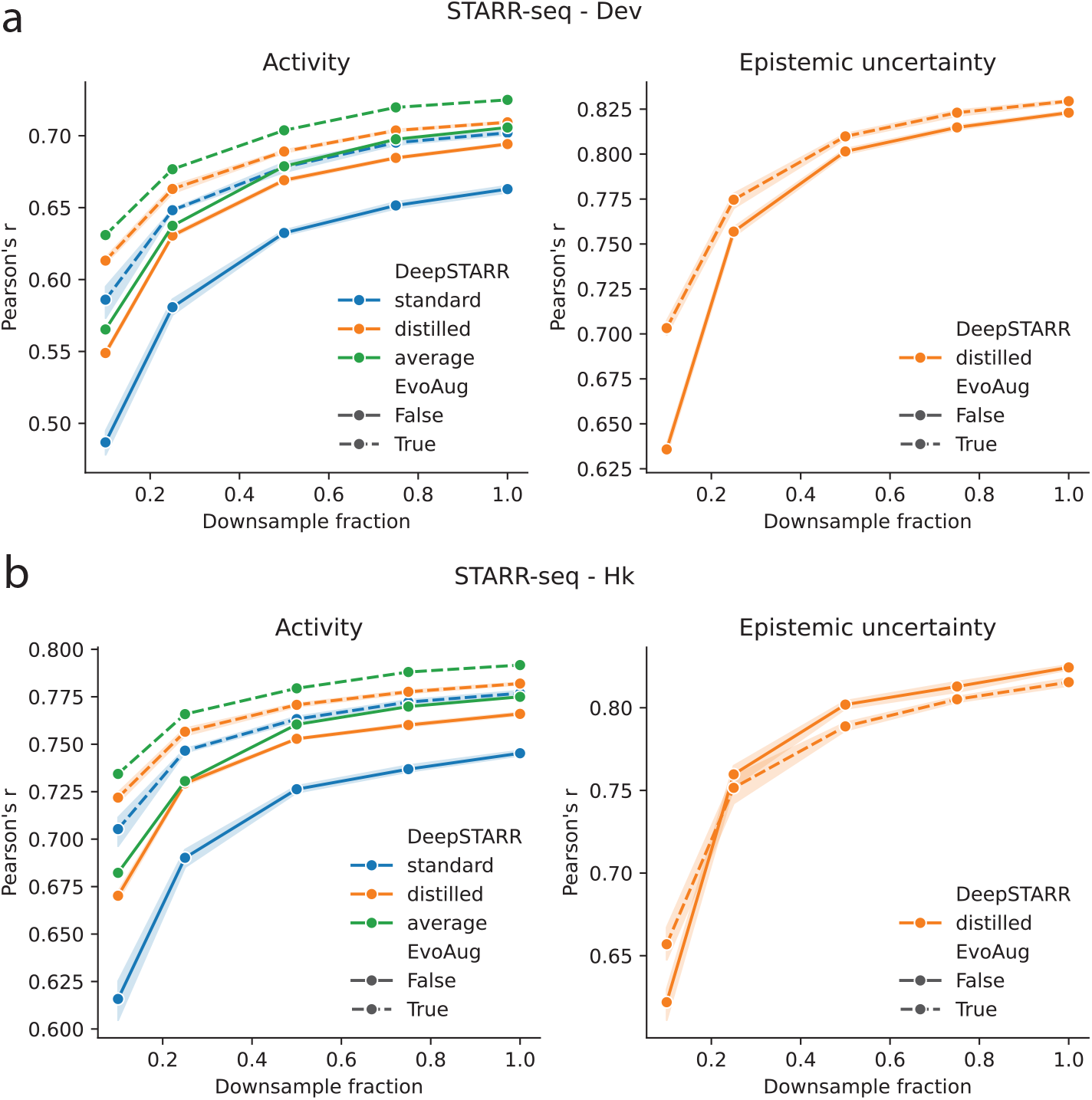
Comparison of model performance with downsampled training data using improved teacher ensembles. (**a**,**b**) Predictive performance for activity (left) and epistemic uncertainty (right) output heads of DeepSTARR models trained on different subsets of randomly downsampled STARR-seq data for (**a**) developmental (Dev) and (**b**) housekeeping (Hk) promoters. Dashed lines indicate DeepSTARR models trained with EvoAug data augmentations, while solid lines represent DeepSTARR models without data augmentations during training. Markers represent the average across 10 models with different random initializations and shaded region indicates 95% confidence interval.

**Supplementary Figure 4.**
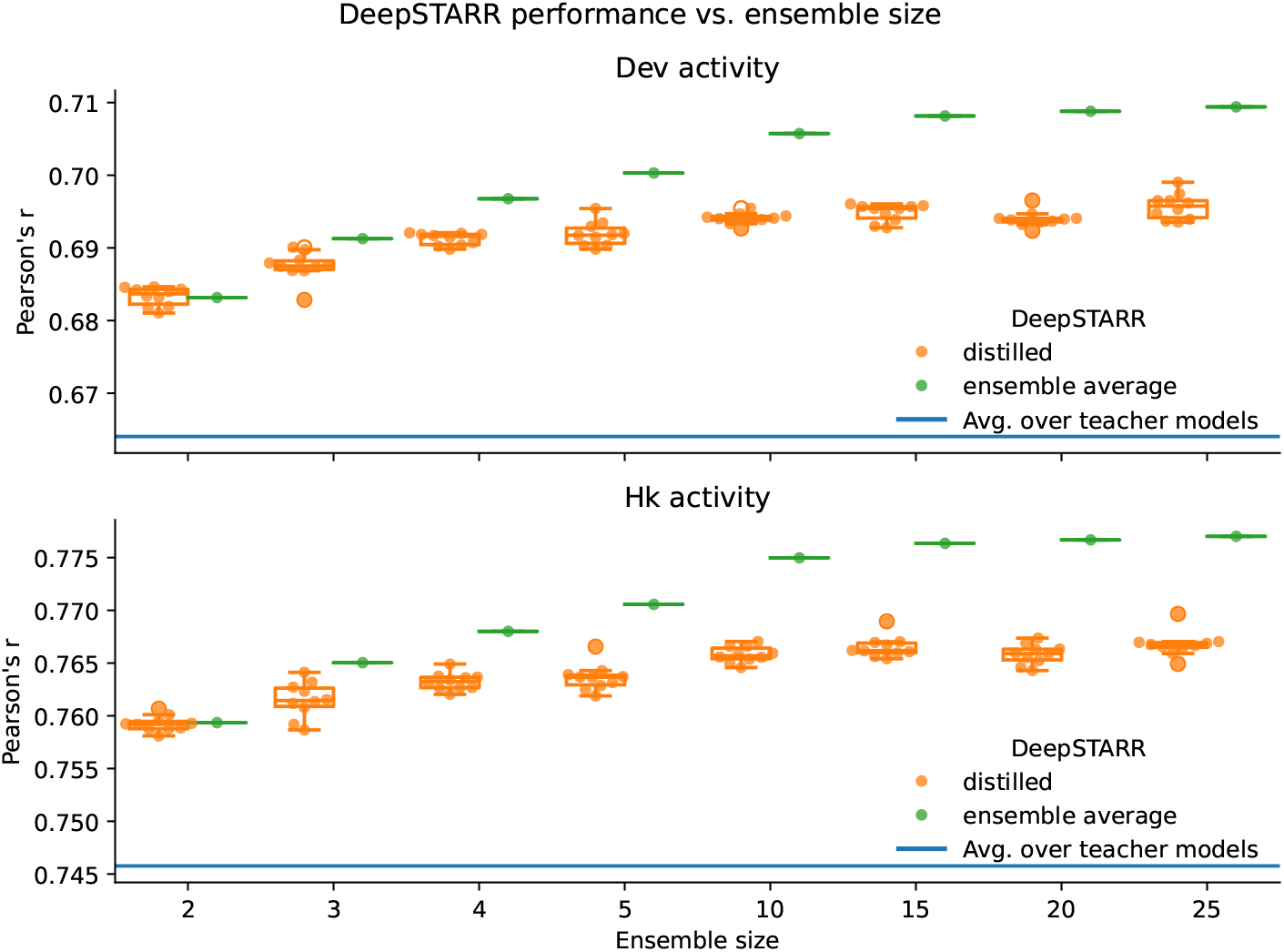
Performance comparison for different teacher ensemble sizes. Boxplot comparison of the performance of DeepSTARR ensembles (green) and distilled DeepSTARR models (orange) for different teacher ensembles comprised of varying number of models. The horizontal blue line represents the average performance metric (Pearson’s *r*) of DeepSTARR with standard training. Boxplots represent *n* = 10 models trained with different random initializations, with the boxes indicating the first and third quartiles, the central line indicates the median, and whiskers denote the data range. Increasing the size of the teacher ensemble yields improvements in predictive accuracy for the ensemble average, but the predictive accuracy of distilled models saturates at a teacher ensemble size of *n* = 10.

**Supplementary Figure 5.**
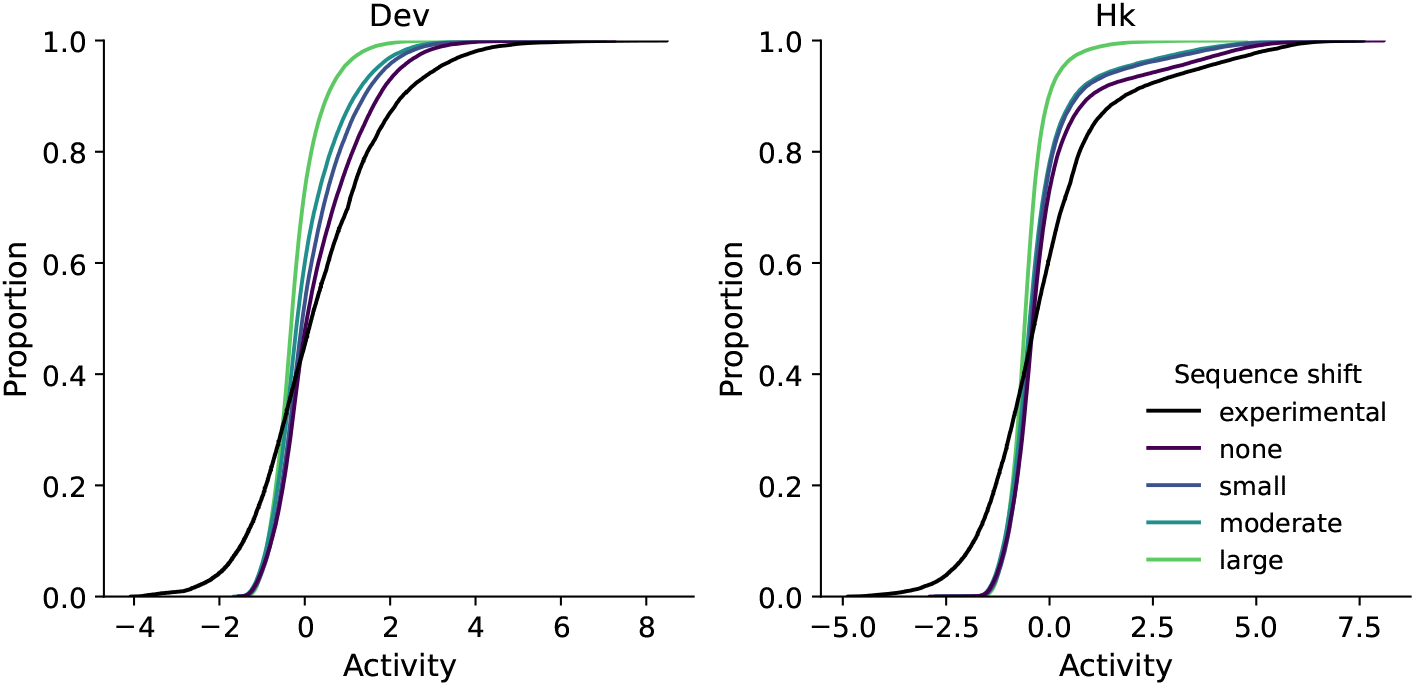
Distribution of predicted activity for STARR-seq test sequences under different distribution shifts. Cumulative distribution plot of predictions given by an exemplary distilled DeepSTARR model for sequences with different degrees of distribution shift: none (original test sequences), small (random mutagenesis with a rate of 5%), moderate (EvoAug mutagenesis), and large (random shuffling).

**Supplementary Figure 6.**
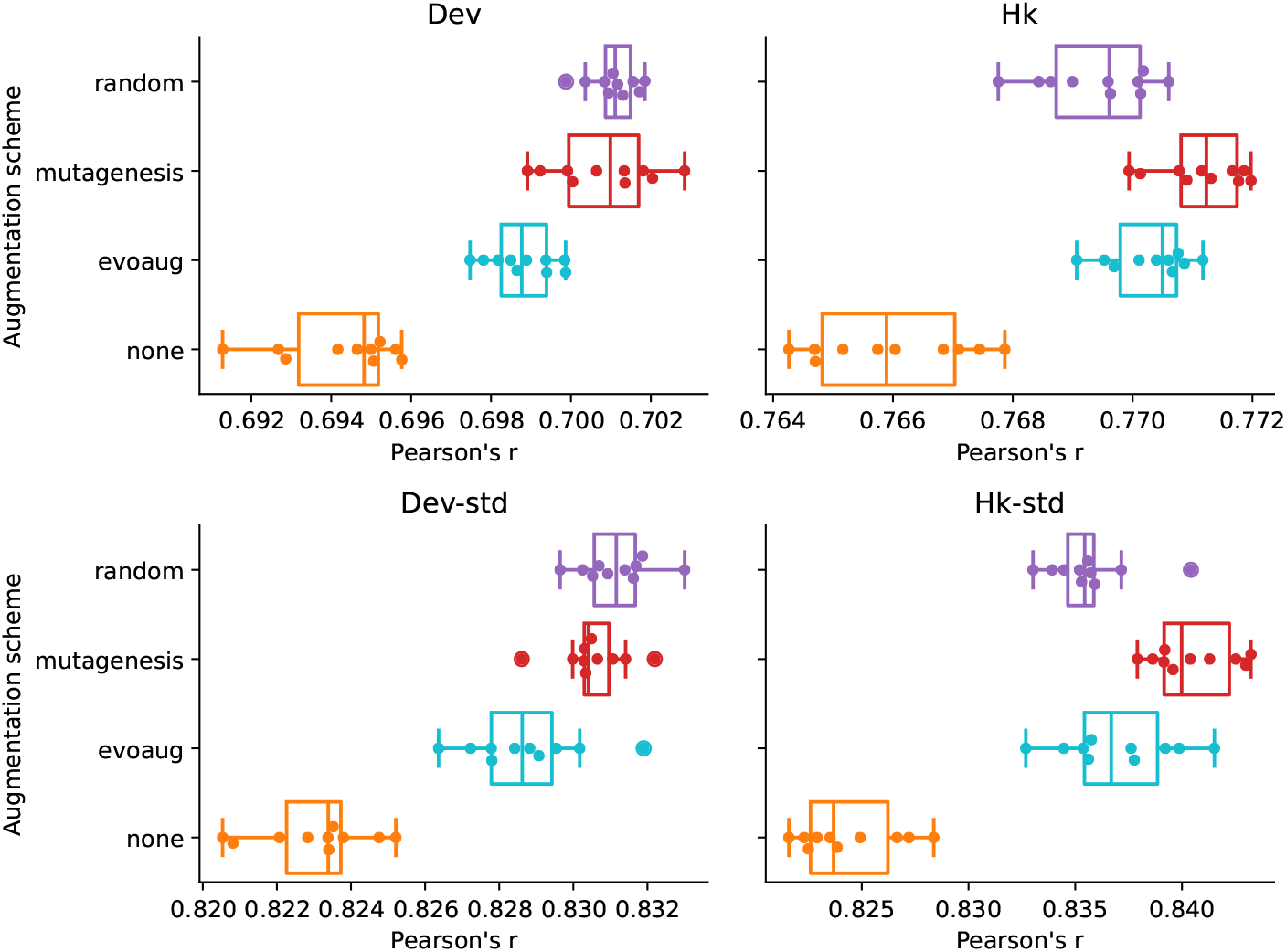
Performance of distilled DeepSTARR models trained with dynamic augmentations. Boxplots of predictive performance of activity head (top) and epistemic uncertainty head (bottom) from distilled DeepSTARR models trained with different dynamic sequence augmentations for Dev (left) and Hk promoters (right). Boxplots represent *n* = 10 models trained with different random initializations, with the boxes indicating the first and third quartiles, the central line indicates the median, and whiskers denote the data range.

**Supplementary Figure 7.**
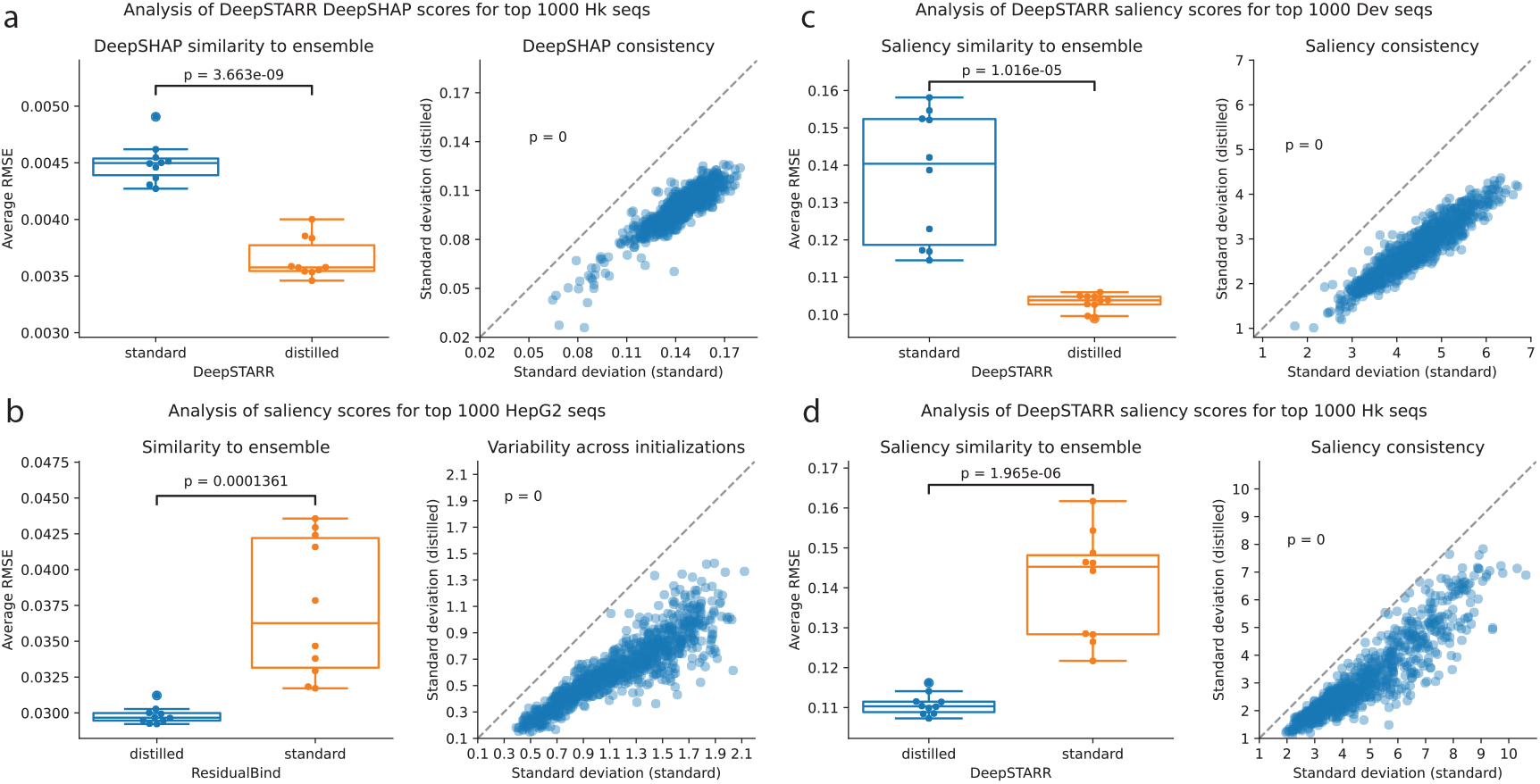
Additional attribution analysis performance comparisons. (left) Boxplot of average root mean squared error (RMSE) between attribution maps generated by individual models trained with standard training (blue) and DEGU distillation (orange), compared to the average attribution map across the teacher ensemble, and (right) scatter plot comparing the standard deviation of attribution scores across individual models trained with standard training (*n* = 10) and DEGU distillation (*n* = 10). Attribution scores were calculated with: (**a**) DeepSHAP for the Hk activity output head of DeepSTARR, (**b**) Saliency Maps for the activity output head of ResidualBind models trained on HepG2 lentiMPRA data, (**c**) Saliency Maps for the Dev activity output head of DeepSTARR, and (**d**) Saliency Maps for the Dev activity output head of DeepSTARR. RMSE is calculated for 1,000 high-activity test sequences. P-values indicate independent two-sided t-tests for average RMSE and paired two-sided t-tests for standard deviation. Boxplots represent *n* = 10 models trained with different random initializations, with the boxes representing the first and third quartiles, the central line indicating the median, and whiskers denoting the data range.

**Supplementary Figure 8.**
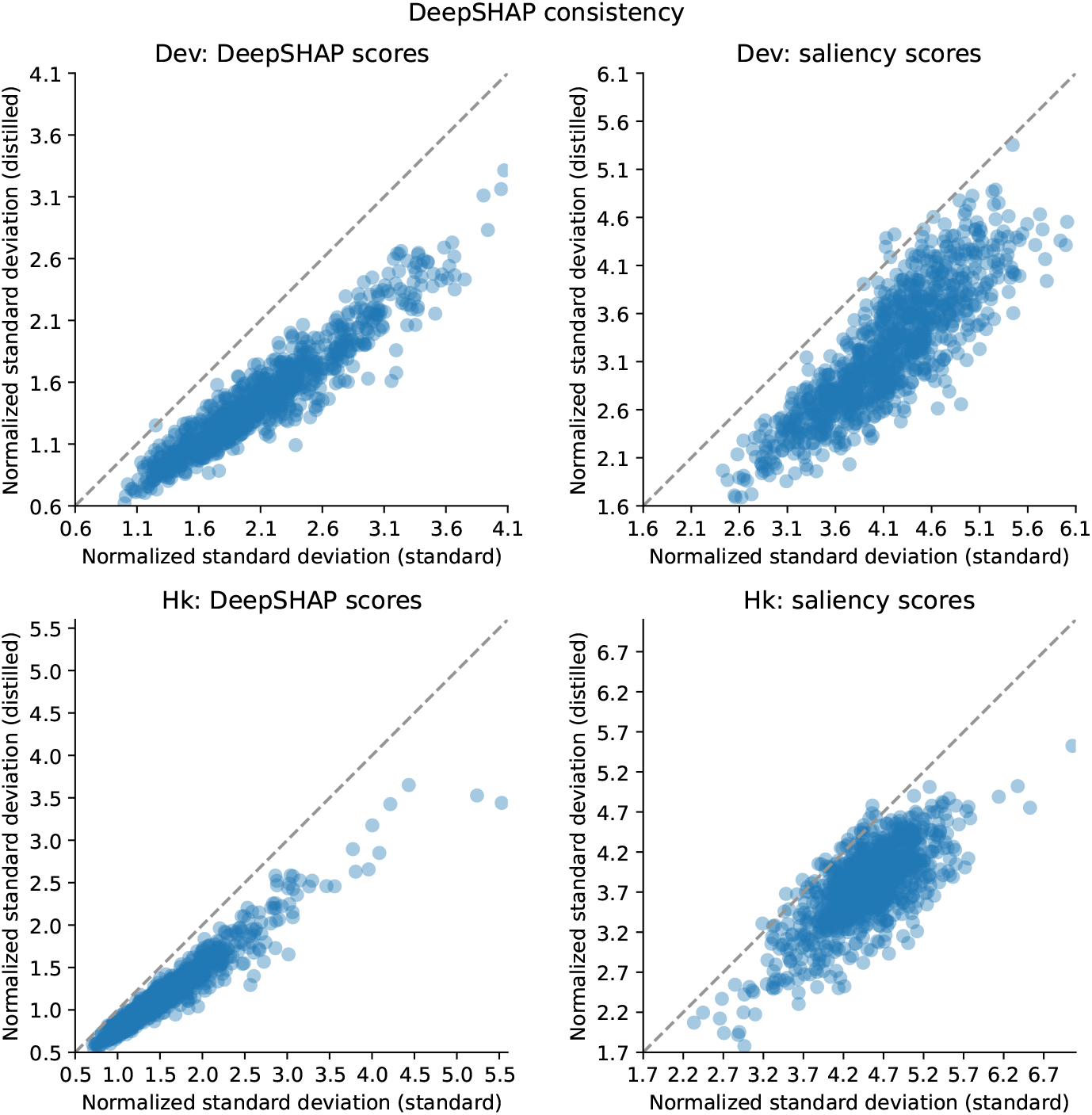
Magnitude-normalized attribution map consistency comparison. Scatter plots comparing the standard deviation of magnitude-normalized attribution scores across individual models trained with standard training (*n* = 10) and DEGU distillation (*n* = 10) for DeepSHAP applied (left) and Saliency Maps (right) applied to DeepSHAP for developmental promoters (top) and housekeeping promoters (bottom). Dots represent *n* = 1, 000 high-activity test sequences.

**Supplementary Figure 9.**
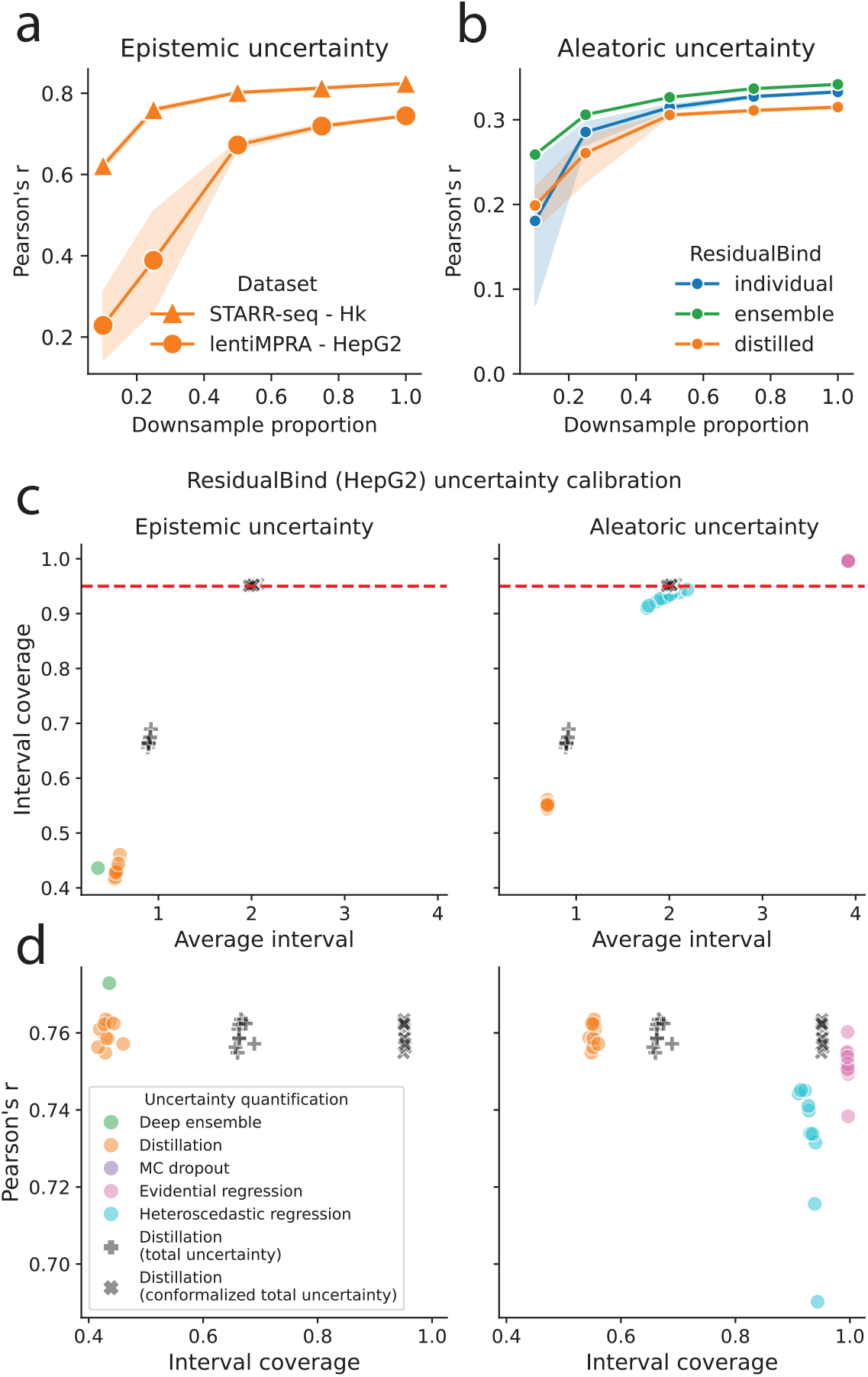
Additional performance comparison of uncertainty estimates. (**a**,**b**) Predictive performance for (**a**) epistemic and (**b**) aleatoric uncertainty for models with standard training (blue), DEGU-distillation (orange), and the teacher ensemble (green), trained on subsets of randomly downsampled training data. Markers represent the average across *n* = 10 models with different random initializations and shaded region indicates 95% confidence interval. Results are shown for (**a**) the Hk epistemic uncertainty head of distilled DeepSTARR models and the epistemic uncertainty output head of distilled ResidualBind models trained on HepG2 lentiMPRA data, and (**b**) the aleatoric uncertainty output heads of ResidualBind models trained on HepG2 lentiMPRA data. (**c**,**d**) Scatter plots of (**c**) prediction interval coverage probability and (**d**) predictive accuracy versus average interval size for different uncertainty quantification methods for epistemic uncertainty (left) and aleatoric uncertainty (right). Uncertainty quantification methods are based on ResidualBind model trained on HepG2 lentiMPRA data. Red dashed line indicates calibration with a 95% interval coverage probability. Each uncertainty quantification method is represented by *n* = 10 dots, indicating a model with different initializations, except for deep ensemble (*n* = 1).

**Supplementary Figure 10.**
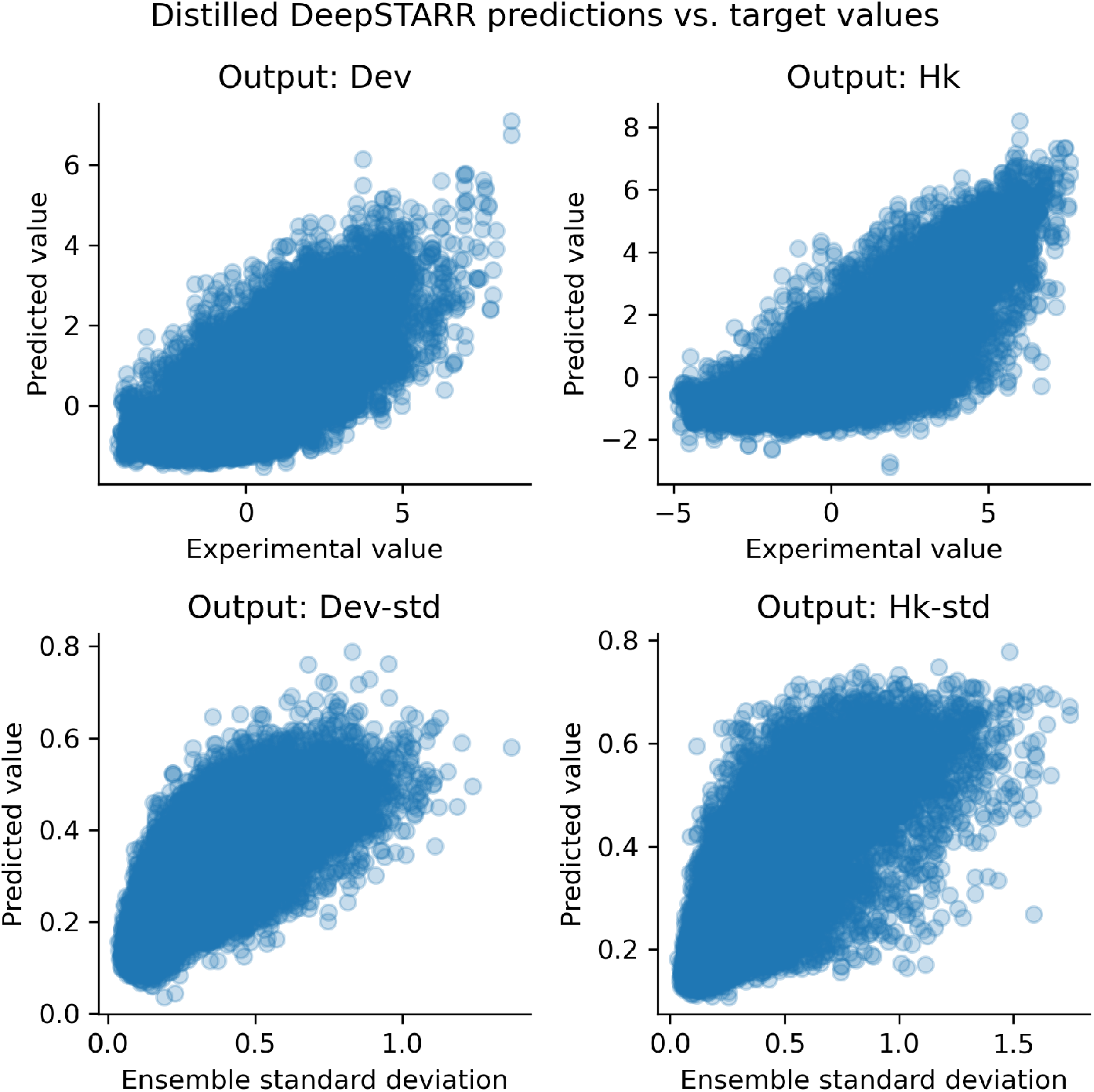
Prediction comparison for distilled DeepSTARR. Scatter plots comparing distilled DeepSTARR model predictions and target values for enhancer activity (top row) and epistemic uncertainty (bottom row) for developmental (left) and housekeeping (right) promoters. Each dot represents a different test sequence (*n* = 41186)

**Supplementary Figure 11.**
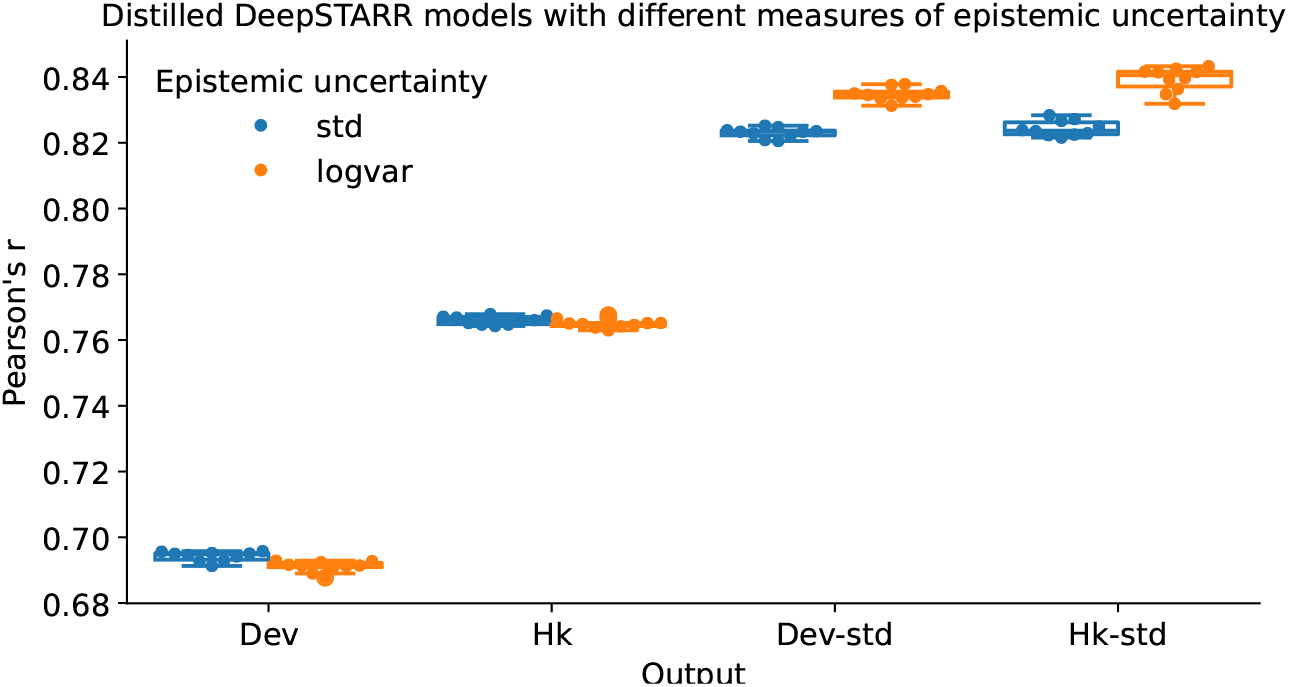
Comparison of different measures of variability. Boxplots of predictive performance for the epistemic uncertainty output head of different distilled DeepSTARR models where epistemic uncertainty ea trained on standard deviation (blue) versus log variance (orange) of the predictions across the teacher ensemble. Boxplots represent *n* = 10 models trained with different random initializations, with the boxes representing the first and third quartiles, the central line indicating the median, and whiskers denoting the data range.

**Supplementary Figure 12.**
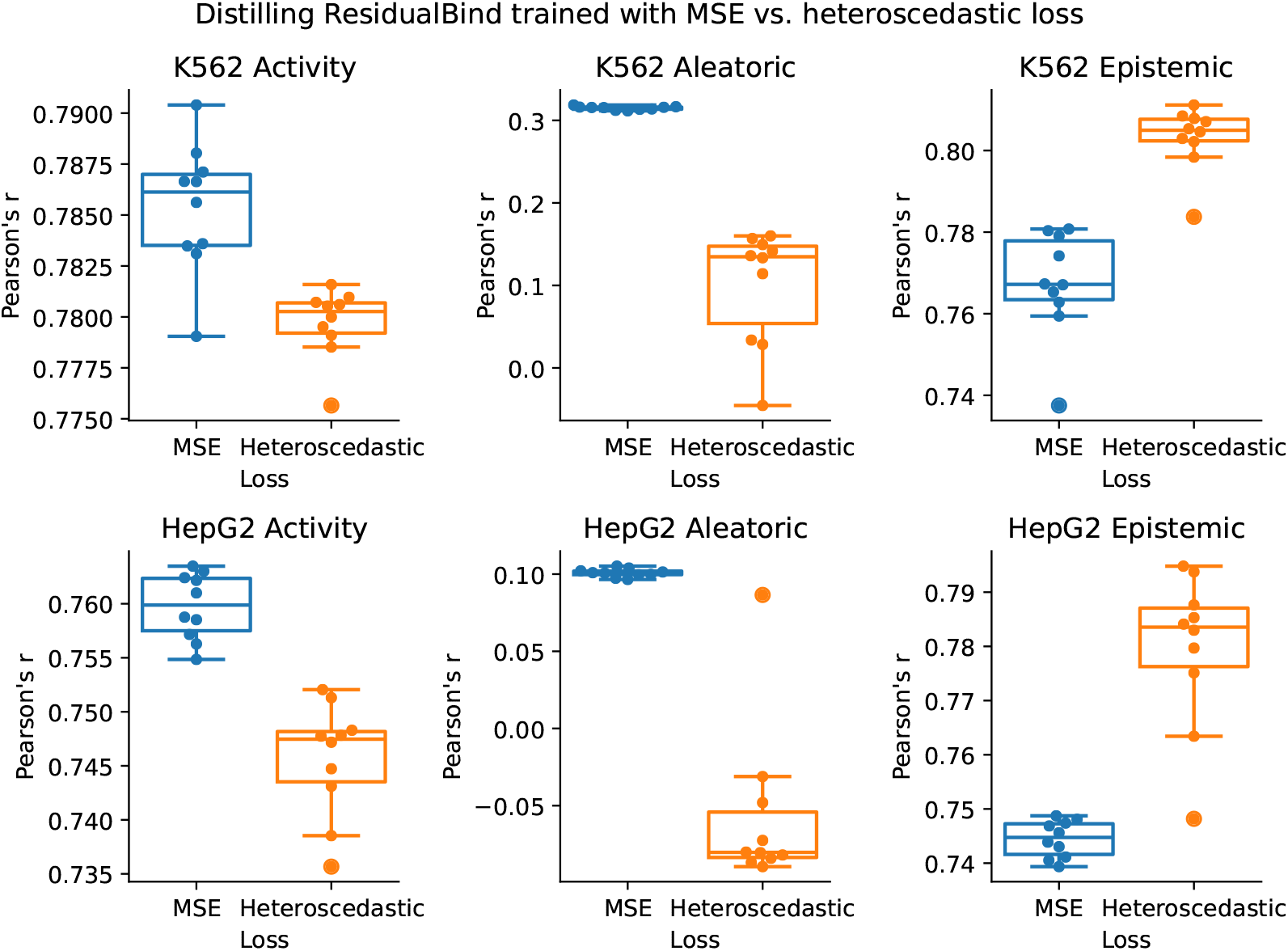
Comparison of loss functions for training ResidualBind. Boxplots comparing performance of ResidualBind trained on lentiMPRA data for K562 (top) and Hepg2 (bottom) using MSE loss or heteroscedastic loss (columns). Models trained with MSE loss learn aleatoric uncertainty from replicate level variation while models trained with heteroscedastic regression implicitly learn aleatoric uncertainty during the training process (but replicate level variation is used to calculate Pearson’s r). Boxplots represent *n* = 10 models with different random initializations, with the boxes representing the first and third quartiles, the central line indicating the median, and whiskers denoting the data range.

**Supplementary Figure 13.**
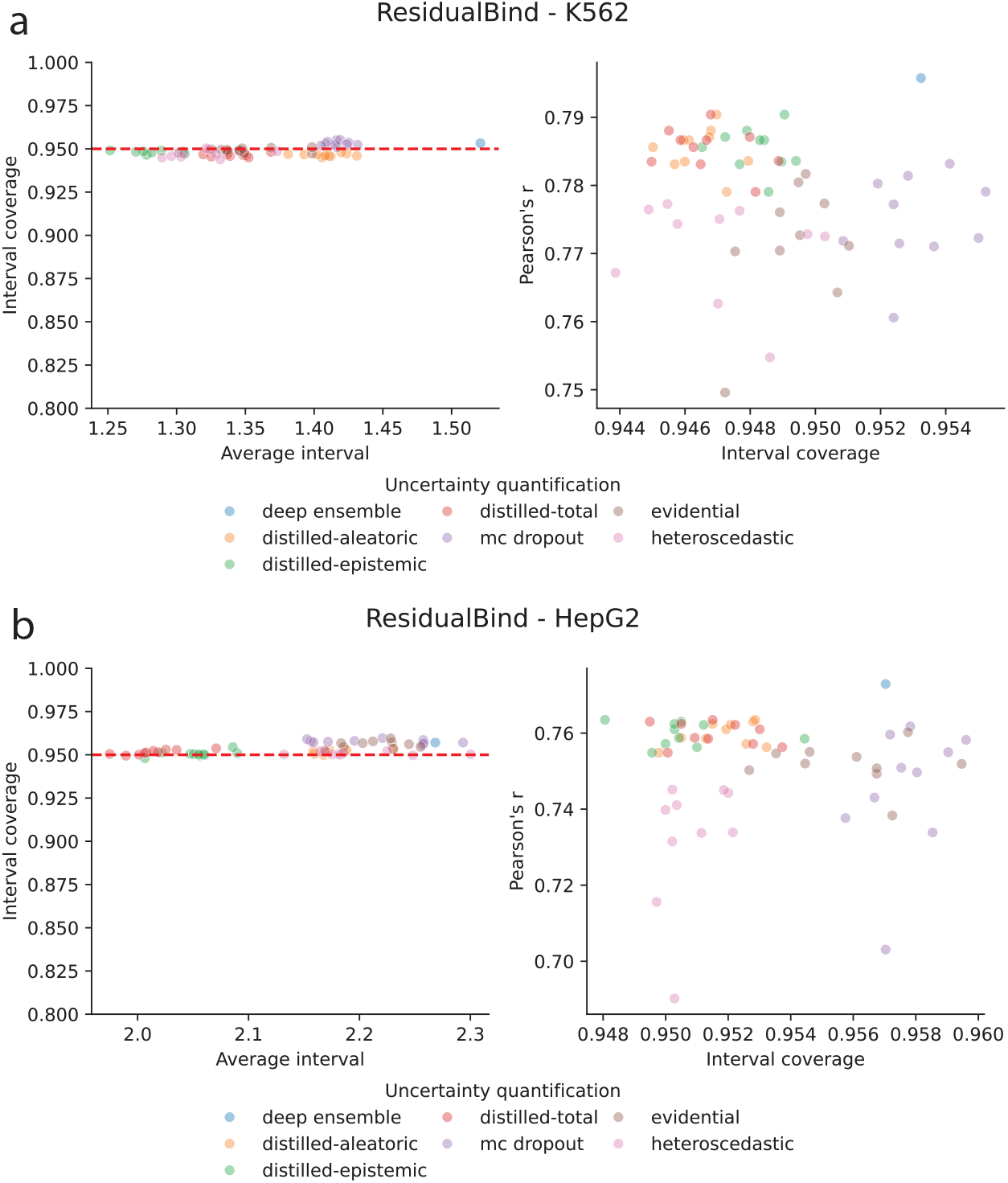
Conformal prediction analysis. Scatter plots of prediction interval coverage probability (left) and predictive accuracy (right) versus average interval size for different uncertainty quantification methods after conformalizing estimates on validation data. The results are shown for ResidualBind models trained on (**a**) K562 and (**b**) HepG2 lentiMPRA data. (left) Red dashed line indicates 95% interval. Each uncertainty quantification method is represented by *n* = 10 dots, indicating a model with different initializations, except for deep ensemble (*n* = 1).

**Supplementary Figure 14.**
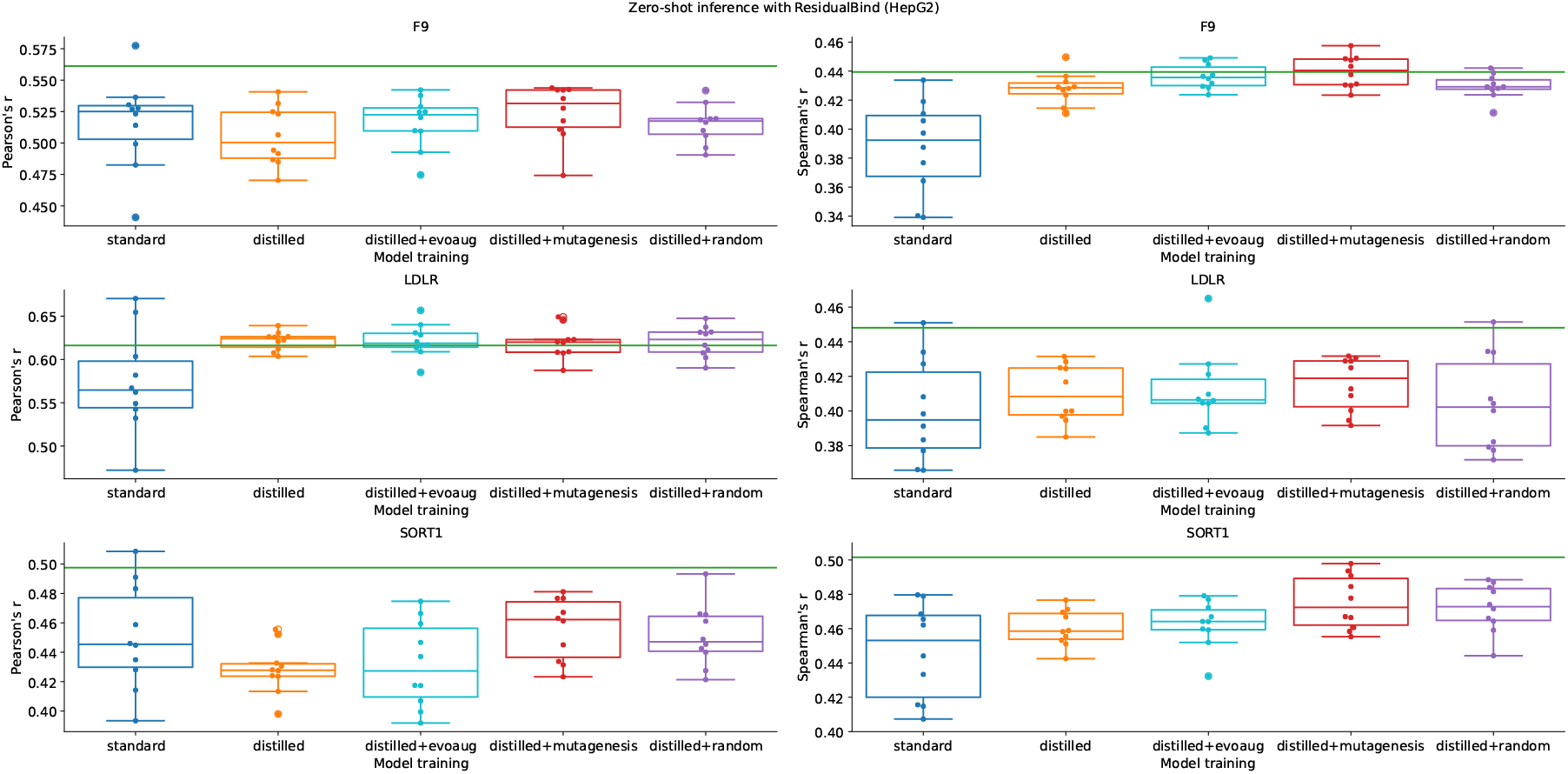
Single nucleotide variant effect prediction for HepG2 regulators. Boxplots of zero-shot variant effect predictive performance for models with standard training (blue); DEGU-distillation (orange); and DEGU-distillation with dynamic augmentations. Boxplots represent *n* = 10 models with different random initializations, with the boxes representing the first and third quartiles, the central line indicating the median, and whiskers denoting the data range. Green horizontal line indicates the performance of the teacher ensemble.

**Supplementary Figure 15.**
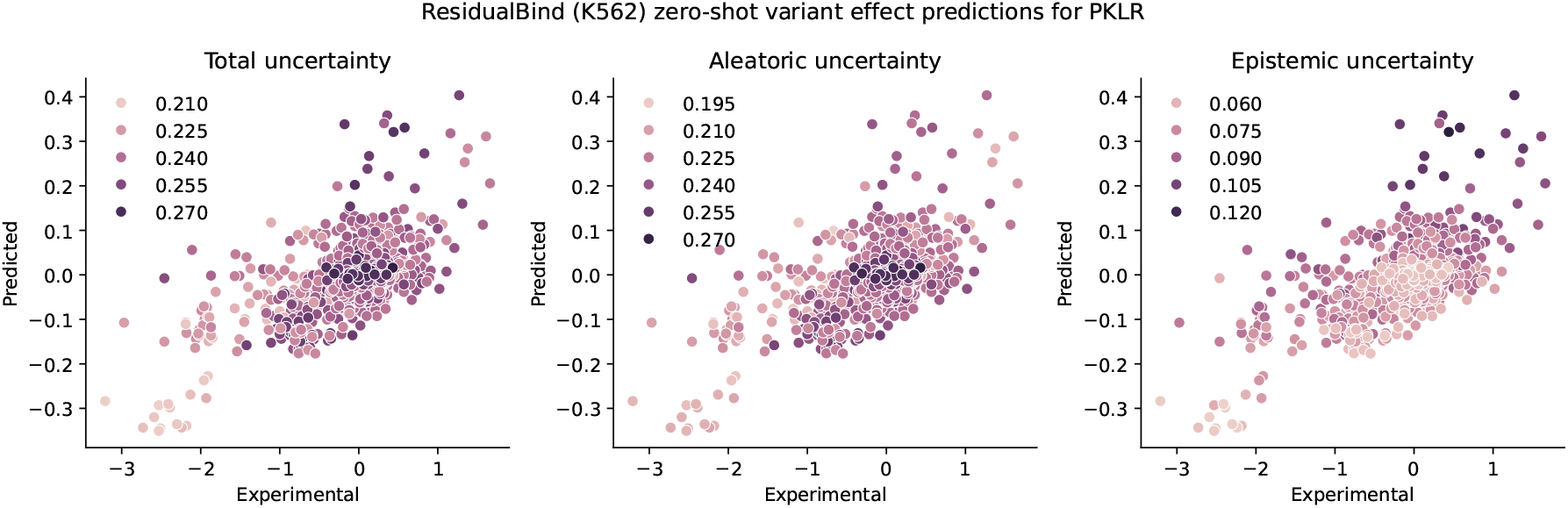
Single-nucleotide variant effect prediction performance with uncertainty annotations. Scatter plots of predictive performance of ResidualBind trained with random mutagenesis augmentations and single-nucleotide variant effects in the PKLR locus measured via an MPRA in K562. The color of each dot represents the uncertainty according to total uncertainty (left), aleatoric uncertainty (middle), and epistemic uncertainty (right).

**Supplementary Figure 16.**
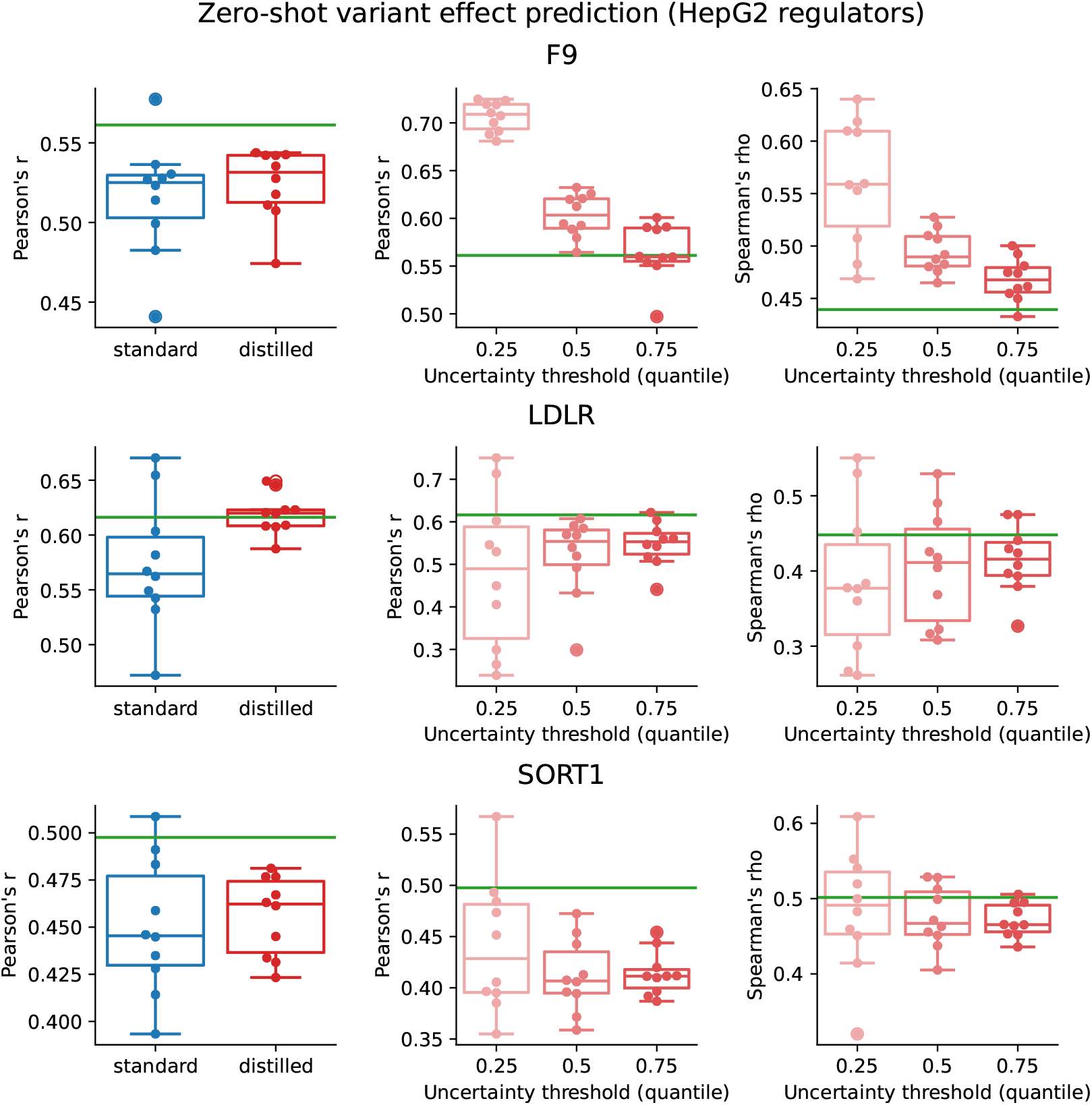
Additional analysis for single-nucleotide variant effect generalization for HepG2 regulators. Boxplots of zero-shot variant effect predictive performance for models with standard training (blue) and DEGU-distillation with mutagenesis augmentations (red) for all nucleotide variants (left) and for variants filtered on different quantile thresholds of predicted total uncertainty (middle, right). Boxplots represent *n* = 10 models with different random initializations, with the boxes representing the first and third quartiles, the central line indicating the median, and whiskers denoting the data range. Green horizontal line indicates the performance of the teacher ensemble.

